# The Australian academic STEMM workplace post-COVID: a picture of disarray

**DOI:** 10.1101/2022.12.06.519378

**Authors:** Katherine Christian, Jo-ann Larkins, Michael R. Doran

## Abstract

In 2019 we surveyed Australian early career researchers (ECRs) working in STEMM (science, technology, engineering, mathematics and medicine). ECRs almost unanimously declared a “love of research”, however, many reported frequent bullying and questionable research practices (QRPs), and that they intended to leave because of poor career stability. We replicated the survey in 2022 to determine the impact of the COVID-19 pandemic and sought more information on bullying and QRPs. Here, we compare data from 2019 (658 respondents) and 2022 (530 respondents), and detail poor professional and research conditions experienced by ECRs. Job satisfaction declined (62% versus 57%), workload concerns increased (48.6% versus 60.6%), more indicated “now is a poor time to commence a research career” (65% versus 76%) from 2019 to 2022, and roughly half reported experiencing bullying. Perhaps conditions could be tolerable if the ecosystem were yielding well-trained scientists and high-quality science. Unfortunately, there are signs of poor supervision and high rates of QRPs. ECRs detailed problems likely worthy of investigation, but few (22.4%) felt that their institute would act on a complaint. We conclude by suggesting strategies for ECR mentorship, training, and workforce considerations intended to maintain research excellence in Australia and improve ECR career stability.

## Introduction

In this paper we discuss survey data collected from Australian early career researchers (ECRs) in 2019 and in 2022, before and after/during the COVID-19 pandemic, with the goal of understanding the pressures on ECRs and on the research community. We focused our study on ECRs working in the STEMM (science, technology, engineering, mathematics and medicine) disciplines in universities and independent research institutes. Our published data from survey of 658 ECRs in 2019, found that ECRs’ ‘love of science’ was their major career motivator, but that most intended to leave their research position because of poor job security [1]. Poor job security is a legitimate problem for Australian researchers. A survey by Professional Scientists Australia in 2021 found that approximately 25% of all science professionals were on a fixed-term contract, and that the average duration was only 18 months [2, 3]. The situation for ECRs in academia is generally worse; our 2019 data suggested that 78% of ECRs were on short-term contracts [1], similar to Hardy et al.’s 2016 report suggesting that >80% of ECRs were on short-term contracts [4].

While competition can drive innovation and productivity, there is always a risk that such pressures will manifest poor or counter-productive behaviour. Our 2019 data suggest that the competitive environment left ECRs vulnerable to exploitation and abuse [1]. Respondents reported alarming rates of bullying and harassment from those in positions of power (31.7% and 25.9% of female and males ECRs, respectively). In depth interviews, conducted in parallel, revealed that ECRs experienced bullying from both senior males and females [5], suggesting a systemic rather than a necessarily gendered problem. Workplace challenges are not unique to the Australian research ecosystem. In 2021, Nature’s international survey data revealed that 27% of respondents had experienced bullying, discrimination, or harassment, with 32% indicating that they had observed such behaviours in their current workplace [6].

The instability associated with short-term contracts is compounded by the subjective nature by which academic output and performance is often quantified. Academic research differs from work in most other sectors of the economy. In the broad economy, most individuals or businesses provide a specific service or product for which the relative value can be quantified. Demand for products/services can be constant, yielding jobs for which fair salary and stability can be anticipated; for example, nurses, teachers or police officers are relatively in constant demand, yielding career stability. By contrast, academic research generally seeks to advance understanding and develop new technologies. Making cutting edge contributions is non-trivial, and it can be challenging to quantify the value of a specific unique contribution as it may take years for the observation/invention to contribute to product development or policy. Because academic outputs are not easily quantifiable, nor easily verifiable in the short-term, it is possible to *game* the so-called metrics. Strategies to game metrics, and create the perception of greater individual productivity, include publication of many [low quality] publications, incorrect allocation of authorship, or, in extreme cases, using fraudulent data to bolster the perceived significance of an individual publication. In our 2019 survey respondents reported an alarming rate of being impacted by questionable research practices (QRPs) at their own institution (41.4% of females and 30.7% of males) [1]. In general, our understanding the impact of QRPs on the scientific community remains limited. While fraud is criminal and likely rare, data suggest that QRPs such as excluding data points may be a prevailing norm, and that these modest but more frequent deviations from truth may have a greater impact on the scientific endeavor [7]. Poor quality or inaccurate data reporting is blamed for the so-called “reproducibility crisis” [8]. A 2015 Nature survey of 1,576 researchers found that 52% agreed that there is a significant crisis of reproducibility, and more than 60% suggested that the cause of poor reproductivity was pressure to publish coupled with selective reporting [8]. Understanding the intertwined nature of QRPs and the precarious employment of ECRs is likely to be essential to understanding if our scientific industry is healthy and legitimately productive.

The plight of ECRs in Australia has been further exacerbated by the COVID-19 pandemic. The Australian university sector is reliant on revenue from international students, which contracted with pandemic-related travel restrictions. In 2020, Universities Australia estimated that Australian universities had lost 17,300 jobs and $1.8 billion in revenue compared to 2019 when we conducted our first survey [1, 9]. A 2022 report suggests that ~35,000 jobs were cut in the university sector, of which ~25% were academic positions (75% were administrative positions) [10]. To understand the impact of the pandemic, we once again surveyed Australian ECRs. We replicated many questions from our 2019 survey, seeking input from January 6 to April 1, 2022. Because of the alarming rate of bullying and harassment and QRPs identified in 2019, the survey was modified to seek additional insight into these workplace problems. To date there has been work on the incidence and impact of bullying and harassment and/or QRPs on ECRs in STEMM fields in other countries [11], but limited work has been performed to understand the research environment in Australia [1, 12, 13].

## Results

### ECR demographics

In this paper we use data from our 2019 survey of 658 Australian ECRs, as previously described in eLife [1]. The new 2022 survey had 530 eligible responses, including 64% who identified as female, 34% as male, and 1% who preferred not to say. Although there were many more women than men this was not unexpected; women were over-represented in our 2019 survey [1]; and it is known that men have lower participation rates in voluntary surveys [14]. The two most common age brackets were 31–35 years old (36.9%) and 36– 40 years old (28.7%), with most respondents having completed their PhD 2–4 years earlier (36.4%) or 5–7 years earlier (31.9%). The four most common countries of birth were Australia (48.9%), England (5.9%), China (3.3%) and India (2.6%). Two thirds (66%) of the respondents held research-only positions at a university or research institute, 25% have combined teaching and research positions. Of the respondents, 48% identified as being in the medical and health sciences. The most recent data from the Australian Research Council (ARC) [15] indicates that 38.9% of Australia’s STEMM workforce is employed in the medical and health sciences (Table 1). Comparison of our survey demographics with this ARC data indicates that our sample and the target population were not statistically different by discipline (*χ*^2^= 11.06 df = 9, p=0.27), and our survey population can be considered representative. A more detailed summary of respondent demographics is provided in Figure 1 (See Supplementary Tables 1 for numerical data shown in Figure 1

**Table 1.** Discipline contribution of Australian STEMM workforce, and discipline contribution to the 2019 and 2022 surveys (2019, n = 658 and 2022, n = 530).

| Discipline | **Percentage of Australian academic STEMM workforce | Percentage of respondents to 2019 survey | Percentage of respondents to 2022 survey |
| --- | --- | --- | --- |
| Mathematical Sciences | 3.8% | 2.8% | 2.6% |
| Physical Sciences | 4.3% | 8.1% | 3.1% |
| Chemical Sciences | 4.7% | 5.7% | 4.2% |
| Earth Sciences | 3.5% | 3.0% | 1.1% |
| Environmental Sciences | 3.2% | 4.0% | 3.0% |
| Biological Sciences | 12.6% | 20.9% | 24.0% |
| Agricultural and Veterinary Sciences | 4.5% | 1.4% | 3.4% |
| Information and Computing Sciences | 6.9% | 2.2% | 4.2% |
| Engineering | 15.4% | 3.6% | 6.0% |
| Technology | 2.1% | 0.8% | 0.9% |
| Medical and Health Sciences | 38.9% | 47.5% | 47.6% |

**Figure 1.**
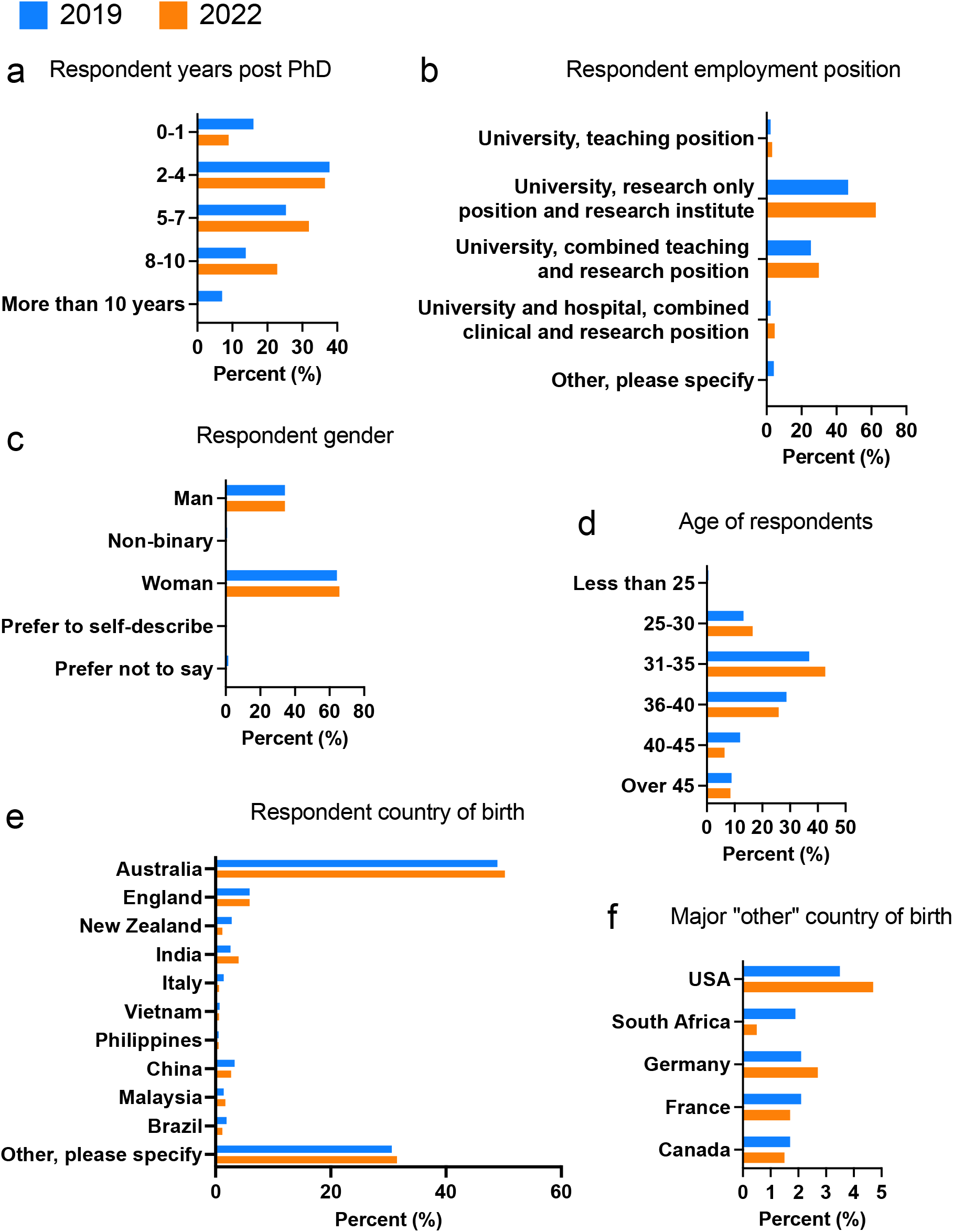
Respondent demographics, **(a)** years post PhD, **(b)** employment contact type, **(c)** respondent gender, **(d)** respondent age, **(e)** respondent country of birth, **(f)** country of birth for respondents selecting the category “other” (2019, n = 658 and 2022, n = 530).

### Workplace culture pre/post pandemic

The undesirable workplace culture identified in 2019 [1] has become less desirable since the COVID-19 pandemic. Overall job satisfaction, already lower than the Australian workforce national average of 80% satisfied [16], has decreased from 62% to 57%. Satisfaction with workplace culture decreased from 51% to 44%. Three-quarters of respondents (76%) agree or strongly agree this is a poor time for a young person to start in this career, compared with 65% in 2019. Similarly, 55% agree or strongly agree that their job is a source of personal strain compared with 52% in 2019.

We examined survey responses by several categories including gender, years postdoctoral, language spoken at home, whether or not the respondent had a disability or chronic health condition, sexual orientation, and research-only versus teaching and research positions. We also compared responses from 2019 with those from 2022, particularly for women, given the reports of the impact of COVID-19 on women [17].

Table 2 shows responses to a range of questions which reflect the workplace culture, and permits comparison of answers before (2019) and after (2022) the pandemic, as well as comparisons by gender. While there are differences between the impact on men and women, the only significant differences in 2022 were for feeling stressed (*χ*^2^= 7.47, df=2, P= 0.024) and inequitable hiring practices (*χ*^2^= 7.51, df=1, P= 0.006) where men were more stressed, but women more frequently reported inequitable hiring practices.

**Table 2.** Workplace culture questions assessed with respect to gender, as well as before and after COVID (2019, n = 658 and 2022, n = 530).

|  | 2019 |  |  | 2022 |  |  |
| --- | --- | --- | --- | --- | --- | --- |
|  | All | Women | Men | All | Women | Men |
| Q13 Overall workload is too high | 48.6% | 52.1% | 42.1% | 60.6% | 61.6% | 60.1% |
| Q15 Overall satisfaction with job (satisfied or very satisfied) | 62.3% | 62.8% | 60.0% | 57.2% | 57.5% | 57.2% |
| Q16-9 Do not feel safe | 12.5% | 11.0% | 15.6% | 13.2% | 14.1% | 11.6% |
| Q16-10 Satisfied with commitment to diversity and inclusiveness | 62.1% | 62.3% | 62.9% | 56.4% | 55.3% | 57.8% |
| Q19 Stressed or v stressed | NA |  |  | 48.4% | 46.4% | 51.8% |
| Q22 Considered leaving for mental health | NA |  |  | 57.5% | 57.4% | 57.6% |
| Q27 Satisfied or very satisfied with workplace culture | 51.0% | 51.9% | 50.3% | 44% | 44.5% | 43.6% |
| Q28 Poor time for a young person to begin – agree & strongly agree | 64.7% | 62.2% | 70.4% | 76.0% | 74.8% | 77.3% |
| Q30-1 inequitable hiring practices | 38.5% | 40.0% | 35.4% | 49.3% | 54.2% | 40.5% |
| Q30-2 Harassment based on power position | 33.5% | 37.1% | 25.9% | 46.3% | 46.5% | 44.8% |
| Q30-3 Impacted by lack of support from institutional superiors | 60.0% | 63.8% | 52.4% | 70.5% | 72.1% | 66.9% |
| Q30-4 Impacted by QRPs in institution sometimes & often | 38.1% | 41.4% | 30.7% | 47.4% | 49.8% | 41.7% |

### Stress and long work hours

As shown in Table 2, 48% of survey respondents reported feeling stressed or very stressed daily. Many (58%) were considering leaving because of because of depression, anxiety, or other mental health concerns related to their work. There is a culture of working long hours in academia [18, 19]. In our 2022 survey, 61% agreed their workload is too high compared with 49% in 2019. Of those who were employed full time and who worked at least 30 hours a week at work, all also worked at home; 20% worked over 16 hours a week at home. Of those who were employed full time and who worked at least 51 hours a week at work, 44% also worked over 11 hours a week at home; 14% worked more than 30 hours a week at home. Lack of work-life balance is a common concern, and this is captured in the comments provided in Table 3.

**Table 3.**
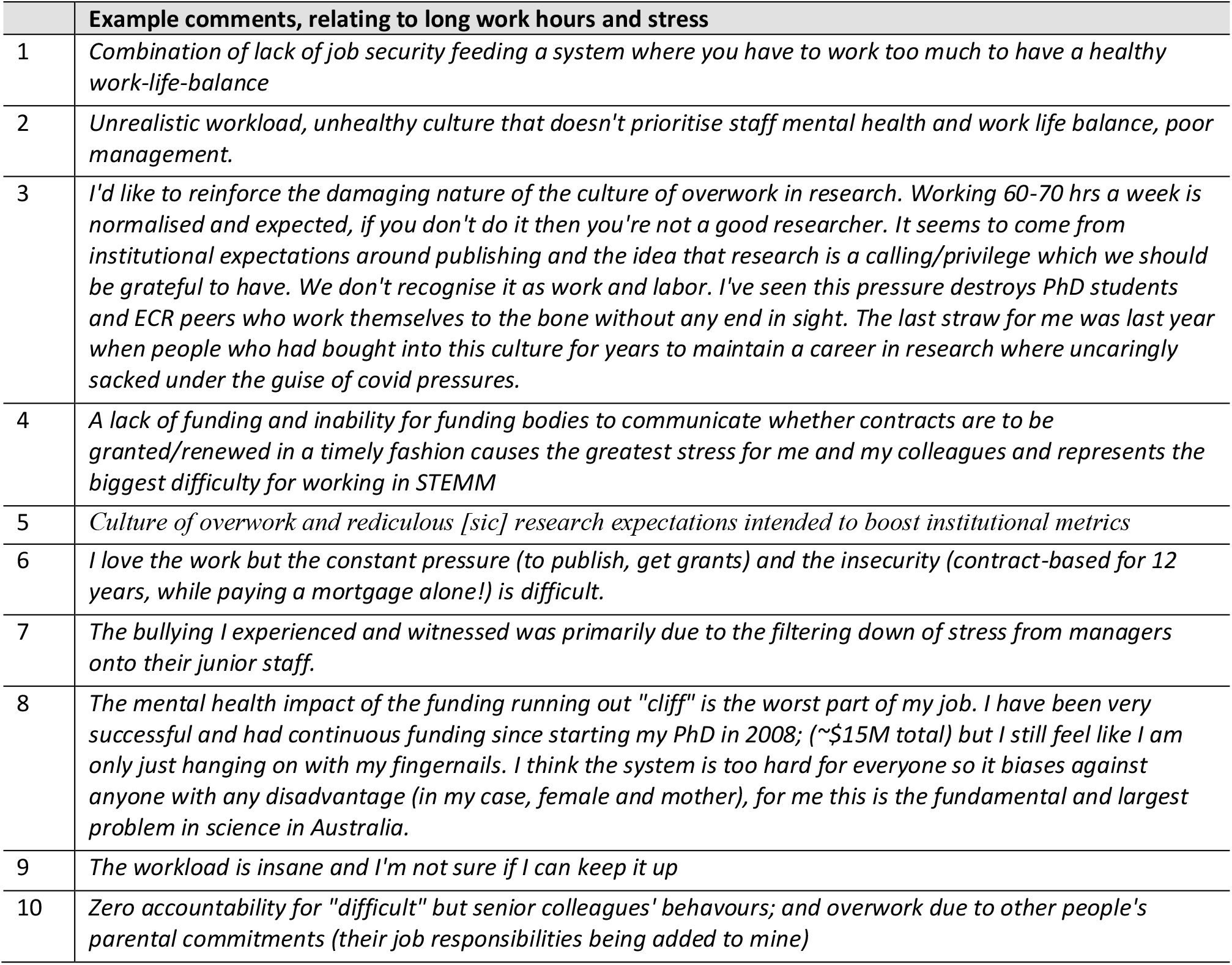
Comments provided in free text answers, selected based on their discussion regarding stress and work hours.

### Intention to leave

The reasons cited for intending to leave academia did not differ greatly between the 2019 and 2022 data (see Table 4). Answers relating to job security, including inadequate job security, lack of funding or lack of independent positions, comprised 89.7% of the reasons ECRs cited for intending to leave research in 2019 and 76.7% in 2022. There were more responses for “other” (14.8%) than in the 2019 survey (8.2%). The principal reasons within “other” related to stress from over work and lack of work life balance, as well as general criticism of the workplace culture, demonstrating that pressures beyond job security are growing.

**Table 4.** Respondent answers to Question 57, “*Why would you leave?*”. (2019, n = 463 and 2022, n = 425).

|  | 2019 |  |  | 2022 |  |  |
| --- | --- | --- | --- | --- | --- | --- |
|  | All | Women | Men | All | Women | Men |
| Inadequate job security | 48.9% | 48.2% | 50.7% | 41.7 % | 38.0 % | 46.5% |
| Lack of funding | 28.8% | 27.8% | 27.6% | 26.1% | 27.4 % | 25.0% |
| Lack of independent positions | 12.0% | 11.6% | 12.7% | 8.9% | 7.7% | 11.8% |
| Family responsibilities | 9.8% | 10.9% | 7.5% | 5.4% | 6.2% | 4.2% |
| Interpersonal problems with your supervisor | 1.3% | 1.3% | 1.4% | 3.1% | 3.7% | 2.1% |
| Other | 8.2% | 9.0% | 6.9% | 14.8% | 17.2% | 10.4% |

### Intention to leave as a function of respondent demographics

Having examined the differences between the answers for men and women we looked for differences when respondents were segmented by years postdoctoral, language spoken at home, disability or chronic health condition, and by sexual preference. Table 5 shows differences according to years postdoctoral. As in 2019, those who were 5-10 years postdoctoral were more likely than those 0-4 years postdoctoral to report that their overall workload was too high, higher impact of harassment based on power position, and that they had experienced bullying. These concerns were reported with a greater frequency in 2022. Conversely, more senior respondents reported a lower level of job satisfaction than their junior colleagues. The differences between groups were only significant for “*overall workload*” (*χ*^2^= 12.92, df=2, P= 0.016).

**Table 5.** Responses grouped by differences by years postdoctoral work.

|  | 2019 |  |  | 2022 |  |  |
| --- | --- | --- | --- | --- | --- | --- |
|  | Years post PhD |  |  | Years post PhD |  |  |
|  | All | 0 to 4 | 5 to 10 | All | 0 to 4 | 5 to 10 |
| Q13 Overall workload is too high | 48.6% | 41.6% | 57.8% | 60.6% | 52.1% | 67.6% |
| Q15 Overall satisfaction with job (satisfied or very satisfied) | 62.3% | 66.9% | 55.7% | 57.0% | 58.7% | 55.6% |
| Q16-9 Do not feel safe | 12.5% | 10.9% | 14.8% | 13.2% | 11.1% | 14.9% |
| Q16-10 Satisfied with commitment to diversity and inclusiveness | 62.1% | 67.2% | 55.2% | 56.4% | 56.5% | 56.4% |
| Q19 Stressed or very stressed | NA |  |  | 48.4% | 46.0% | 50.4% |
| Q22 Considered leaving for mental health | NA |  |  | 57.5 % | 58.0 % | 57.0 % |
| Q27 Satisfied or very satisfied with workplace culture | 51.0% | 54.0% | 46.8% | 44.0% | 48.1% | 40.5% |
| Q28 Poor time for a young person to begin – agree and strongly agree | 64.7% | 58.5% | 73.2% | 76.0% | 73.1% | 78.4% |
| Q30-1 inequitable hiring practices | 38.5% | 35.8% | 41.3% | 49.3% | 48.6% | 49.8% |
| Q30-2 Harassment based on power position | 33.5% | 31.1% | 36.7% | 46.3% | 41.5% | 50.2% |
| Q30-3 Lack of support from institutional superiors | 60.0% | 54.1% | 67.5% | 70.5% | 67.5% | 73.0% |
| Q30-4 Impacted by QRPs in institution sometimes and often | 38.1% | 35.8% | 41.3% | 47.4% | 46.2% | 48.3% |

We tabulated results based on country of birth (see Supplementary Table 2), finding that those not born in Australia were more often concerned about inequitable hiring practices (55.9% versus 40.9% Australian born). There was broad concern from ECRs on this issue, with concerns increasing across all ECRs demographics from 38.5% in 2019 to 49.3% in 2022. We considered that contrasting those born or not born in Australia might not capture challenges faced by those from different cultural or language backgrounds. To explore this possibility, we assessed respondent data as a function of whether the ECR spoke English at home (see Table 6). The most remarkable difference in the data for 2022 is the increased spread in the views about whether “*it is a poor time for a young person to begin a research career*” between those who speak English at home and those who do not. English speakers more often indicated that now is not good time to begin a research career than the non-English speakers (83.4% versus 61.7% difference in 2022 is significant, *χ*^2^ = 18.7, df=2, p<0.001). We presume that those who do not speak English at home either immigrated or live with family that have immigrated to Australia; note that only five individuals who do not speak English at home reported Australia as their country of birth. Those who spoke English as their first language more often cited mental health as a reason for intending to leave (60% versus 51.7%). Counterintuitively, while those who do not speak English at home appeared to be less pessimistic, they also appeared to be less satisfied with the workplace culture (*χ*^2^= 9, df=2, P= 0.011). Perhaps those from another country valued a job in Australia more than one in their country of origin. Overall, language appeared to only play a minor role in ECRs’ outlooks on their work environment, and it is concerning that for most of these workplace challenges approximately 50% of all respondents expressed concern.

**Table 6.**
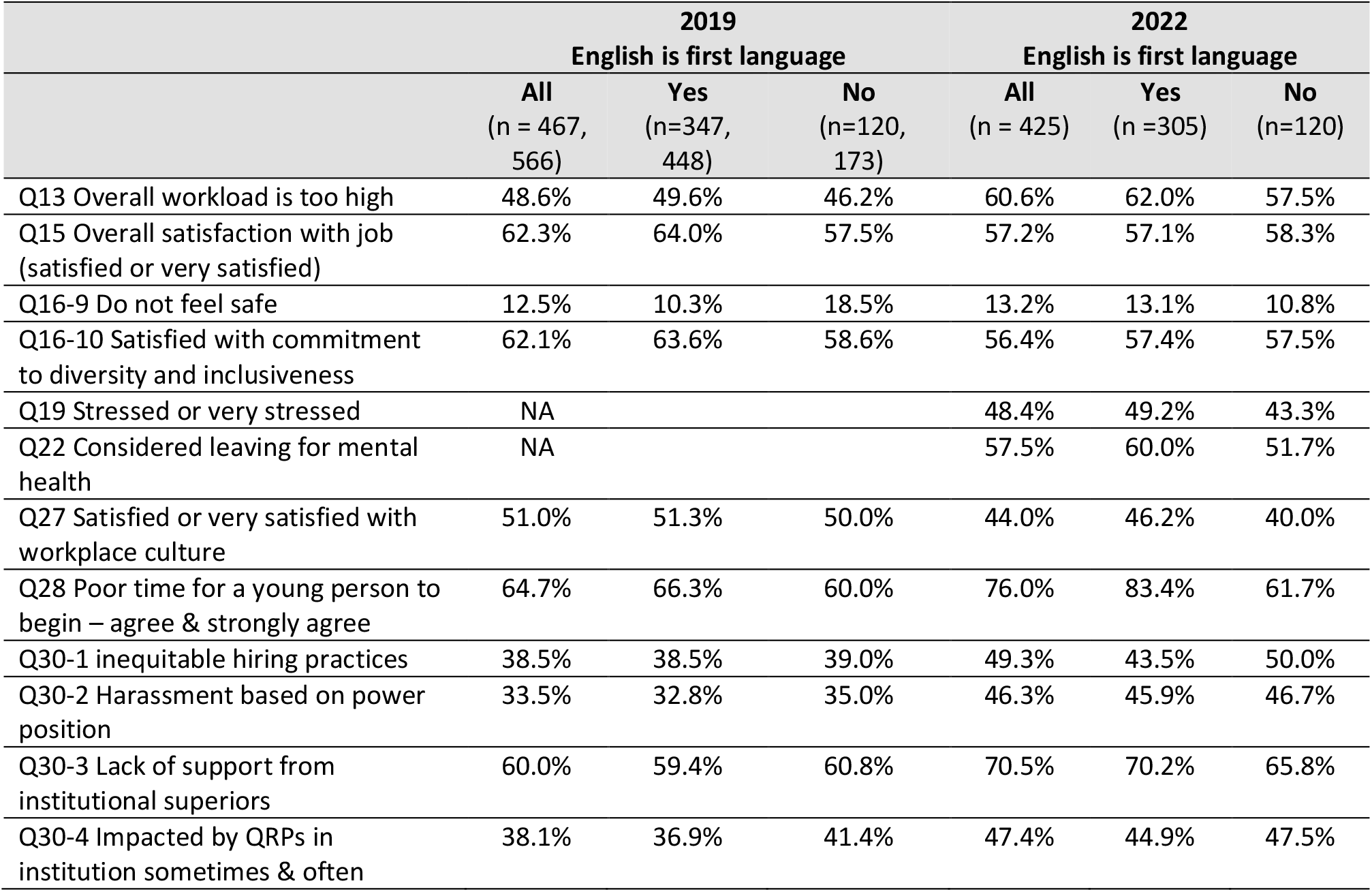
Responses grouped by differences by language spoken at home.

|  | 2019 |  |  | 2022 |  |  |
| --- | --- | --- | --- | --- | --- | --- |
|  | English is first language |  |  | English is first language |  |  |
|  | All<br>(n = 467,<br>566) | Yes<br>(n=347,<br>448) | No<br>(n=120,<br>173) | All<br>(n = 425) | Yes<br>(n =305) | No<br>(n=120) |
| Q13 Overall workload is too high | 48.6% | 49.6% | 46.2% | 60.6% | 62.0% | 57.5% |
| Q15 Overall satisfaction with job<br>(satisfied or very satisfied) | 62.3% | 64.0% | 57.5% | 57.2% | 57.1% | 58.3% |
| Q16-9 Do not feel safe | 12.5% | 10.3% | 18.5% | 13.2% | 13.1% | 10.8% |
| Q16-10 Satisfied with commitment<br>to diversity and inclusiveness | 62.1% | 63.6% | 58.6% | 56.4% | 57.4% | 57.5% |
| Q19 Stressed or very stressed | NA |  |  | 48.4% | 49.2% | 43.3% |
| Q22 Considered leaving for mental<br>health | NA |  |  | 57.5% | 60.0% | 51.7% |
| Q27 Satisfied or very satisfied with<br>workplace culture | 51.0% | 51.3% | 50.0% | 44.0% | 46.2% | 40.0% |
| Q28 Poor time for a young person to<br>begin – agree & strongly agree | 64.7% | 66.3% | 60.0% | 76.0% | 83.4% | 61.7% |
| Q30-1 inequitable hiring practices | 38.5% | 38.5% | 39.0% | 49.3% | 43.5% | 50.0% |
| Q30-2 Harassment based on power<br>position | 33.5% | 32.8% | 35.0% | 46.3% | 45.9% | 46.7% |
| Q30-3 Lack of support from<br>institutional superiors | 60.0% | 59.4% | 60.8% | 70.5% | 70.2% | 65.8% |
| Q30-4 Impacted by QRPs in<br>institution sometimes & often | 38.1% | 36.9% | 41.4% | 47.4% | 44.9% | 47.5% |

It is common for PhD candidates and ECRs to travel internationally for education or work [20], and often individuals pursue immigration in their host nation. It is worth considering how employment stability impacts the immigration process, and the additional stressor this may have on foreign born STEMM ECRs in Australia. Table 7 includes quotes from respondents who highlight this specific challenge.

**Table 7.** Comments provided in free text answers on the topic of immigration.

|  | Example comments, relating to immigration |
| --- | --- |
| 1 | *It often feels like if you want to stay in Australia after your PhD then academia is the only option* |
| 2 | *I had to work for the uni for 5 years to get them to write me a letter for PR [permanent residency]. They kept me on 1 year contracts so contracts were "too short to offer that"* |
| 3 | *I had no choice. I needed a job within the field to be able to apply for PR.* |
| 4 | *Short term contracts are particularly stressful for immigrants on a working visa as it makes it virtually impossible to apply for PR or change job* |
| 5 | *Australia with its hard VISA conditions poses an additional threat to the constant job-renewals.* |

In Table 8 we investigated the impact of a disability or chronic health condition on ECRs. We did not collect these data in 2019, but the fact that 119 of 530 respondents (22.5%) identified as having a disability or chronic health condition demonstrates the importance of capturing these data. It is useful to know that 15% of all respondents indicated that they face barriers or limitations in their day-to-day activities because of chronic health issues or disabilities. Of those who reported disability or chronic health condition, 65% say they have mental health issues; 10% say they have a visual disability, 8% report they have dyslexia and 6% say they have a hearing disability. A further 10% say they have a listed disability but would prefer not to say what it is. For this cohort, respondents who suffer a disability more frequently considered leaving for mental health reasons (73.1% versus 52.8%), reported not feeling safe (19.3% versus 13.2%), were stressed (53.8% versus 48.4%), had been harassed based on power position (51.4% versus 46.3%), experienced bullying (66.7% versus 47.8%), or had been pressured regarding authorship (57.8% versus 49.0%). The differences between people with and without a disability are statistically significant for “*do not feel safe*” (*χ*^2^= 6, df=2, p=0.05), “*considered leaving for mental health*” (*χ*^2^=13, df=2, p=0.002) and “*experienced bullying*” (*χ*^2^=24, df=2, p<0.001). Some might argue that a disability could make individuals pre-disposed or vulnerable to such challenges, but when 65% of subgroup, or 77 out of the 530 total respondents, report suffering mental health disability it may be worth considering the impact of the work environment. Similarly, it is worth noting that those who do not cite a disability nor mental health problems also report high levels of concern across all categories.

**Table 8.** Responses grouped by disability or chronic health condition. Those that responded “*prefer not to say*” (n = 13) to the disability question were grouped with those reporting a disability for the analysis presented in this table.

|  | 2022 |  |  |
| --- | --- | --- | --- |
|  | All<br>(n =496) | No<br>Disability<br>(n=350) | Disability<br>(n=132) |
| Q13 Overall workload is too high | 60.6% | 61.9% | 56.3% |
| Q15 Overall satisfaction with job (Satisfied or very satisfied) | 57.2% | 55.7% | 62.2% |
| Q16-9 Do not feel safe | 13.2% | 11.1% | 19.3% |
| Q16-10 Satisfied with commitment to diversity and inclusiveness | 56.4% | 57.3% | 54.6% |
| Q19 Stressed or very stressed | 48.4% | 46.2% | 53.8% |
| Q22 Considered leaving for mental health | 57.5% | 52.8% | 73.1% |
| Q27 Satisfied or v satisfied with workplace culture | 44% | 45.4% | 40.5% |
| Q28 Poor time for a young person to begin – agree & strongly agree | 76.0% | 77.1% | 72.1% |
| Q30-1 inequitable hiring practices | 49.3% | 50.0% | 46.9 % |
| Q30-2 Harassment based on power position | 46.3% | 43.7% | 51.4% |
| Q30-3 Lack of support from institutional superiors | 70.5% | 68.9% | 73.9% |
| Q30-4 Impacted by QRPs in institution sometimes & often | 47.4% | 44.9% | 53.2% |
| Q33 Experienced bullying | 47.8% | 41.1% | 66.7% |
| Q41-1 Pressure regarding authorship | 49.0% | 46.3% | 57.8% |

Next, we analyzed respondent data as a function of sexual orientation (see Table 9). Of those respondents who declared their sexual orientation, 16.6% self-declared as LGBTIQ. There was little difference between responses from people who reported they were heterosexual compared with those who said they were LGBTIQ. However, LGBTIQ respondents reported higher rates of being impacted by QRPs (57.1% versus 43.7%, (*χ*^2^=6.1, df=2, p=0.014) relative to their heterosexual peers. Although not quite statistically significant, LGBTIQ people also reported higher rates of inequitable hiring practices (57.1% versus 46.7%, *χ*^2^=3.66, df=1, p=0.056).

**Table 9.** Responses grouped by sexual orientation (those who indicated that they “prefer not to say” on the gender question were not included in the analysis shown in this table).

|  | 2022 |  |  |
| --- | --- | --- | --- |
|  | All | Heterosexual<br>(n = 332) | LGBTIQ<br>(n = 66) |
| Q13 Overall workload is too high | 60.6% | 60.8% | 59.6% |
| Q15 Overall satisfaction with job (satisfied or very satisfied) | 57.2% | 59.0% | 55.3% |
| Q16-9 Do not feel safe | 13.2% | 12.7% | 13.5% |
| Q16-10 Satisfied with commitment to diversity and inclusiveness | 56.4% | 59.0% | 50.4% |
| Q19 Stressed or very stressed | 48.4% | 46.7% | 51.1% |
| Q22 Considered leaving for mental health | 57.5% | 57.8% | 60.0% |
| Q27 Satisfied or very satisfied with workplace culture | 44.0% | 44.0% | 46.4% |
| Q28 Poor time for a young person to begin – agree & strongly agree | 76.0% | 75.6% | 78.6% |
| Q30-1 inequitable hiring practices within institution | 49.3% | 46.7% | 57.1% |
| Q30-2 Harassment based on power position | 46.3% | 46.7% | 50.9% |
| Q30-3 Lack of support from institutional superiors | 70.5% | 69.6% | 73.3% |
| Q30-4 Impacted by QRPs in institution sometimes & often | 47.4% | 43.7% | 57.1% |

### Influence of contract type

We compared respondents who were employed in research-only positions versus teaching and research positions (see Table 10). In Australia, most on-going, tenure-equivalent, positions are teaching and research appointments. Researchers appreciate that an additional teaching load can limit research productivity but often pursue teaching as a strategy to achieve career stability. This reasoning is consistent with reported workload concerns, with 58% of research-only respondents indicating that their workload was too high, compared with 72% for those on teaching and research contracts (*χ*2= 7.28, df=2, P= 0.026). Although not significantly different, inequitable hiring practices appear to trouble those on teaching and research positions more than those in research-only positions. It is possible that those exerting additional effort to teach feel especially aggrieved if they are unsuccessful in realizing contract renewal or advancement.

**Table 10:** Responses grouped by research-only versus teaching and research position.

|  | 2019 |  |  | 2022 |  |  |
| --- | --- | --- | --- | --- | --- | --- |
|  | All | Research only | Teaching and Research | All | Research only | Teaching and Research |
| Q13 Overall workload is too high | 49.2% | 41.4% | 64.4% | 60.6% | 57.9% | 71.3% |
| Q15 Overall satisfaction with job (satisfied or very satisfied) | 63.4% | 64.5% | 59.1% | 57.0% | 57.7% | 55.9% |
| Q16-9 Do not feel safe | 12.9% | 12.6% | 14.1 % | 13.2% | 12.3% | 15.8% |
| Q16-10 Satisfied with commitment to diversity and inclusiveness | 63.0% | 63.8% | 60.7 % | 56.4% | 56.5% | 55.1% |
| Q19 Stressed or very stressed | NA | NA | NA | 48.4% | 49.9% | 45.6% |
| Q22 Considered leaving for mental health | NA | NA | NA | 57.7% | 57.7% | 62.4% |
| Q27 Satisfied or v satisfied with workplace culture | 51.3% | 53.2% | 45.6 % | 44.0% | 46.2% | 37.6% |
| Q28 Poor time for a young person to begin – agree & strongly agree | 65.5% | 66.3% | 67.2 % | 76.0% | 76.7% | 76.9 % |
| Q30-1 inequitable hiring practices | 38.6% | 33.4% | 46.8% | 49.3% | 46.2% | 55.6% |
| Q30-2 Harassment based on power position | 33.8% | 31.7% | 41.3% | 46.3% | 44.9% | 47.0% |
| Q30-3 Lack of support from institutional superiors | 58.3% | 54.5% | 64.3 % | 70.5% | 73.5% | 73.5% |
| Q30-4 Impacted by QRPs in institution sometimes & often | 36.0% | 37.3 % | 34.4% | 47.4% | 47.2% | 53.0% |

Next, we analyzed respondent data as a function of continuing versus short-term contracts (see Table 11). The only significant difference between the groups was concern that workloads were too great (68.7% for continuing positions, 56.4% for contract positions of 1-3 years, and 61.5% for contract positions < 1 year; (*χ*^2=^16.19, df=4 p=0.003). For all three groups, concern about high workload was greater than in 2019, prior to the pandemic. We expected that those having on-going positions would view many of the challenges in academia more favourably and were surprised to find that ECRs had similar concerns regardless of contract stability. Perhaps most striking is that regardless of contract stability, approximately half (44.7% to 50.8%) had experienced harassment from someone in a position of power, nearly three quarters expressed concern about lack of support from institutional superiors (68.5% to 74.0%), and approximately half claim to have been impacted by QRPs at their own institution sometimes or often (46.0% to 53.0%). Across all three areas, the situation has degraded substantially from 2019, with averages in harassment from a someone in a position of power increasing from 33.5% to 46.3%, lack of support from institutional superiors increasing from 60.0% to 70.5%, and the frequency by which ECRs were impacted by QRPs at their own institute increasing from 38.1% to 47.4%. The impact of the COVID-19 pandemic on researchers in Australia [21] and internationally [22, 23] and its consequent impact on work challenges is also highlighted by example free text comments provided in Table 12.

**Table 11:** Responses grouped by continuing versus short-term contracts.

|  | 2019 |  |  |  | 2022 |  |  |  |
| --- | --- | --- | --- | --- | --- | --- | --- | --- |
|  | All | Continuing<br>Position | Contract<br>position<br>1-3 years | Contract<br>position<br><1 year | All | Continuing<br>Position<br>n=147 | Contract<br>position<br>1-3 years<br>n=259 | Contract<br>position<br><1 year<br>n=52 |
| Q13 Overall workload is too high | 48.6% | 62.6% | 42.9% | 46.9% | 60.6% | 68.7% | 56.4% | 61.5% |
| Q15 Overall satisfaction with job (satisfied or very satisfied) | 62.3% | 63.1% | 63.9% | 51.6% | 57.2% | 61.3% | 58.3% | 43.1% |
| Q16-9 Do not feel safe | 12.5% | 10.3% | 12.0% | 19.2% | 13.2% | 14.8% | 12.7% | 15.7% |
| Q16-10 Satisfied with commitment to diversity and inclusiveness | 62.1% | 57.7% | 64.0% | 53.8% | 56.4% | 50.7% | 58.7% | 56.9% |
| Q19 Stressed or very stressed | NA | NA | NA | NA | 48.4% | 44.3% | 49.6% | 52.9% |
| Q22 Considered leaving for mental health | NA | NA | NA | NA | 57.5% | 55.7% | 60.0% | 51.0% |
| Q27 Satisfied or very satisfied with workplace culture | 51.0% | 44.8% | 52.0% | 44.9% | 44.0% | 36.4% | 48.5% | 40.0% |
| Q28 Poor time for a young person to begin – agree & strongly agree | 64.7% | 56.0% | 64.3% | 78.1% | 76.0% | 72.0% | 77.5% | 84.0% |
| Q30-1 inequitable hiring practices | 38.5% | 40.4% | 31.5% | 56.9% | 49.3% | 54.5% | 46.0% | 50.0% |
| Q30-2 Harassment based on power position | 33.5% | 39.3% | 29.7% | 38.9% | 46.3% | 50.8% | 44.7% | 46.0% |
| Q30-3 Lack of support from institutional superiors | 60.0% | 70.8% | 53.5% | 68.1% | 70.5% | 73.5% | 68.5% | 74.0% |
| Q30-4 Impacted by QRPs in institution sometimes & often | 38.1% | 33.7% | 38.1% | 48.6% | 47.4% | 53.0% | 46.4% | 46.0% |

**Table 12.** Comments from free text about how the COVID-19 pandemic has impacted workload.

|  | **Example comments, relating to the influence of COVID-19 on workload** |
| --- | --- |
| 1 | *COVID has increased exponentially the workload.* |
| 2 | *It is my direct lab supervisor who gives no encouragement and always keeps my position 'hanging' with short term contracts, no opportunities to develop my skills or move up in my career. This was made worse during home schooling due to covid lockdowns* |
| 3 | *The transition to online teaching during the pandemic provided a huge burden for academics. I live in a state with lock-downs and working from home being enforced. Research had to be put on hold for two years, and yet the university does not seem to be acknowledging this in workload agreements for 2021 and 2022.* |
| 4 | *The area I am in is female dominated (not male) however the males within the area tend to get promoted faster, get offered other opportunities others aren't aware of, have greater leniencies with children at home than women do, etc. COVID has increased exponentially the workload.* |
| 5 | *The last straw for me was last year when people who had bought into this culture for years to maintain a career in research where uncaringly sacked under the guise of covid pressures.* |
| 6 | *I my case across covid our institute has been pretty supportive and on most points generally pretty good, however, these view were not carried through my supervisor. As far as she is concern through out the pandemic it should be business as usual and there is no reason why you should just work harder to make up for lost time* |
| 7 | *The transition to online teaching during the pandemic provided a huge burden for academics. I live in a state with lock-downs and working from home being enforced. Research had to be put on hold for two years, and yet the university does not seem to be acknowledging this in workload agreements for 2021 and 2022. If anything, the workload agreements have been skewed so that less time is given for teaching and administration even though online classes take much more administrative work and have to be re-designed from any existing face-to-face format.* |
| 8 | *The enforced isolation of lockdowns and working from home has made it difficult to connect to the community when starting at new institutions. I've never met or spoken to the majority of my current colleagues.* |
| 9 | *But now there are no jobs to apply for as Universities have doubled down for the covid winter, so I must continue to exist with one foot in science and one foot out. My family and I deserve more financial security.* |
| 10 | *The job losses in the university sector, and instability of research-only positions (one - three year contracts, rather than ongoing positions ) means that as an ECR, I'm "stuck" in my current position and feel like I have no bargaining power when it comes to negotiating research and teaching loads* |

### Mentorship and supervisor guidance

Academia is a challenging career. While the magnitude of the concerns communicated above may be surprising, most in the industry would have some awareness of these challenges because they are increasingly being aired in major reports or journals [11, 24, 25]. We asked ECRs about the support they have received from their supervisor(s) over the previous 12 months (see Table 13). Despite the known challenges for ECRS and career stability, only 63.2% of supervisors discussed career aspirations with ECRs, 24.7% discussed skill development, 11.0% discussed alternative career options, and 9.1% of supervisors did not engage any of the items listed in Table 13 over the past 12 months. These data do not have to be seen as an indictment of supervisors; instead, it could be used as a reminder that these questions are all important, and that asking these questions may spontaneously initiate mentorship activities. Supervisors are not able to modify the national research ecosystem, and many are likely struggling to maintain their own employment or career progression, but they may be able to impart wisdom and provide support to help ECRs to at least ask these important questions. Table 14 provides comments from respondents who describe challenges with their supervision; these comments highlight pressures on supervisors, as well as instances where supervisors likely have not behaved appropriately.

**Table 13.** ECRs describing support from their supervisors in response to the following questions (Q53; *Has your supervisor, PI or manager done any of the following within the last 12 months?)*

| Answer |  |
| --- | --- |
| Had a conversation with you about your career aspirations? | 63.2% |
| Provided career advice and guidance? | 54.5% |
| Discussed your performance? | 63.6% |
| Provided an example of appropriate ethical codes? | 9.6% |
| Noted your achievements? | 52.2% |
| Offered you training to support your skill development? | 24.7% |
| Provided an example of appropriate research standards? | 13.1% |
| Connected you to others within or outside your field? | 37.3% |
| Supported you with personal issues? | 28.9% |
| Supported your wellbeing? | 38.9% |
| Provided expert advice? | 43.1% |
| Conducted a formal appraisal? | 28.9% |
| Discussed alternative career options? | 11.0% |
| Requested your feedback on their management of you? | 7.2% |
| None of the above? | 9.1% |
| Not applicable? | 0.9% |

**Table 14.**
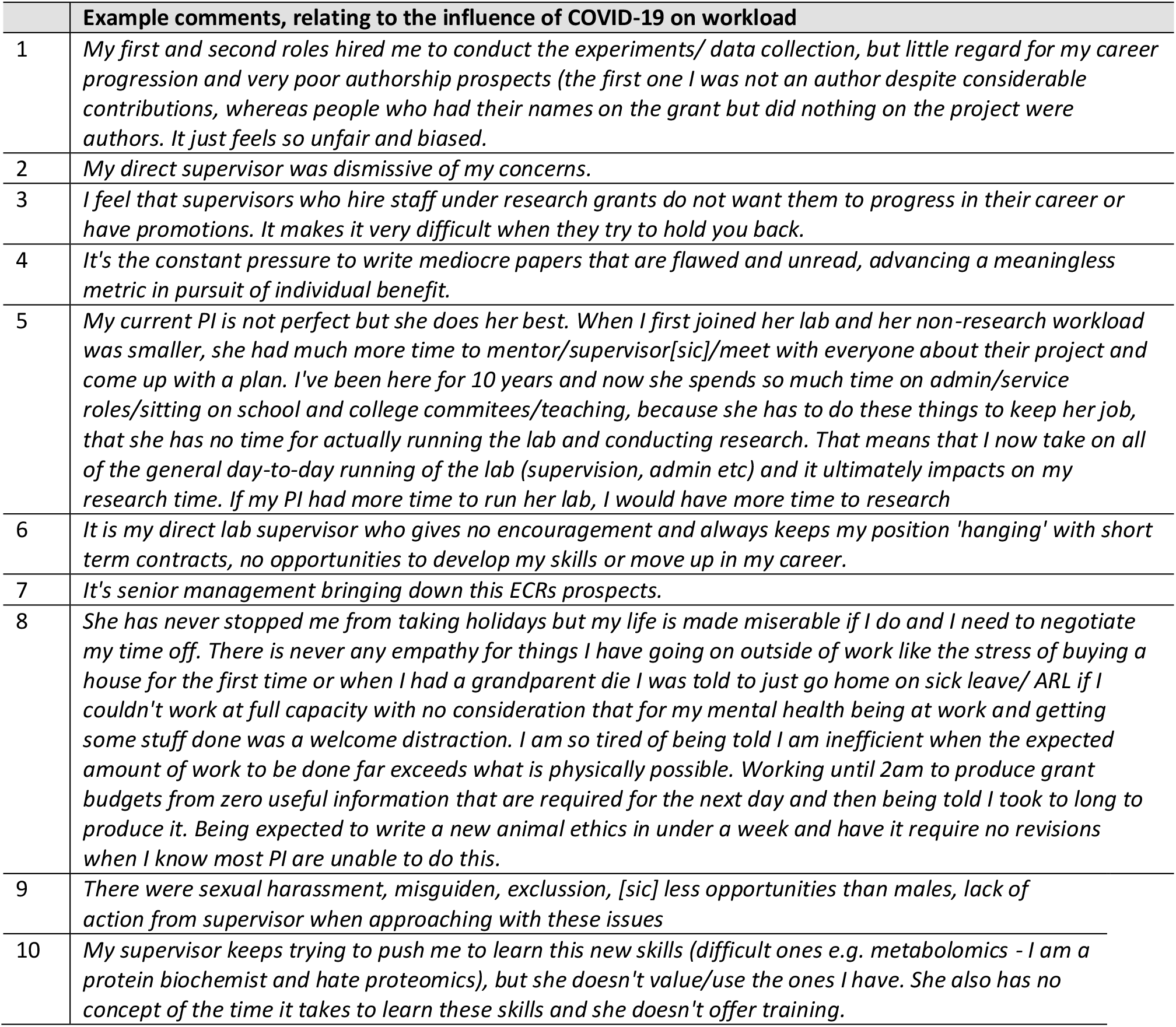
ECRs describing weak support from their supervisors.

### Bullying, harassment, and discrimination

Respondents to our 2019 survey reported a high prevalence of bullying and harassment and QRPs [1], however that survey did not provide respondents with an opportunity to offer a full explanation. In the 2022 survey we investigated both bullying and harassment and QRPs in greater depth. Table 15 provides detail as to who was observed to be harassing or bullying from the perspective of respondents who experienced the event(s), or respondents who had observed the event(s). For those who experienced the event, 42.5% were perpetrated by the supervisor, 31.3% by another senior colleague, or 21.1% by a peer, respectively. For those who observed the event(s), 42.6% were perpetrated by the supervisor, 44.0% by another senior colleague, or 20.9% by a peer, respectively. These numbers are reasonably consistent, suggesting that supervisors are more often the perpetrators, although rates of abuse from senior colleagues and peers are equally concerning. It may be that the similar frequency indicates either certain common behaviours are viewed as bullying or harassment, or that these behaviours are common because they are tolerated by both supervisors and senior colleagues, and to a lesser extent by peers. We sought to determine if there appeared to be a causal basis for harassment or bullying (see Table 16)

**Table 15.** For those who experienced or observed bullying and harassment, who was the perpetrator?

| All | Bullying or harassing perpetrator |  |  |  |  |
| --- | --- | --- | --- | --- | --- |
|  | Supervisor | Other senior colleague | Peer | Prefer not to say | Other |
| Of the 47.8% (n = 339) of those who experienced harassment or bullying: Who was the perpetrator? (select all that apply) | 42.5% | 31.3% | 21.1% | 4.1% | 0% |
| Of the 65.5% (n = 474) of who observed harassment or bullying: Who was the perpetrator? (select all that apply) | 42.6% | 44.0% | 20.9% | 2.5% | 0% |

**Table 16.** What appeared to be the basis of the harassment or bullying?

| All | Basis of the harassment or bullying |  |  |  |  |
| --- | --- | --- | --- | --- | --- |
|  | Gender | Race or ethnicity | Age | Nationality | Class/socio-economic background |
| Of the 47.8% (n = 339) of those who experienced harassment or bullying, what was the bullying or discrimination experienced behaviour related to? | 22.9% | 9.7% | 11.0% | 7.7% | 5.0% |
| Of the 65.5% (n = 474) of who observed harassment or bullying, what was the bullying or discrimination behaviour witnessed related to? | 21.6% | 15.8% | 9.3% | 11.1% | 5.2% |

In our 2022 survey data (Table 16), of those who experienced harassment or bullying, 22.9% felt it had been motivated by gender, 9.7% by race/ethnicity, 11.0% by age, 7.7% by nationality, and 5.0% by class or socio-economic background. For those who observed harassment or bullying, 21.6% felt that it had been motivated by gender, 15.8% by race/ethnicity, 9.3% by age, 11.1% by nationality, and 5.2% by class or socioeconomic background. The numbers for those experiencing and observing these behaviours are similar, again perhaps indicating that certain behaviours are commonly repeated in workplaces.

It is likely one of the reasons that bullying and discrimination continues to be problematic is that ECRs have relatively low confidence that institutional leaders will take their concerns seriously. For example, when respondents were asked if they felt that concerns related to their experiences of bullying or discrimination would be listened to, 55.7% said yes, 22% said no, and 22.3% were unsure (n = 463). Worse, when asked if respondents felt their concerns regarding bullying or harassment would be acted upon, only 26.4% said yes, 36.5% said no, and 37.2% were unsure (n = 463). Finally, when asked if respondents would feel comfortable speaking out about instances of bullying or discrimination without suffering negative personal consequences, only 36.1% said yes, 31.3% said no, 30.9% were unsure, and 1.7% preferred not to say (n = 463). Finally, example comments from those who experienced or observed bullying or harassment are provided in Table 17 and 18. These comments provide perspective on the type and severity of bullying, and the impact on individuals.

**Table 17.**
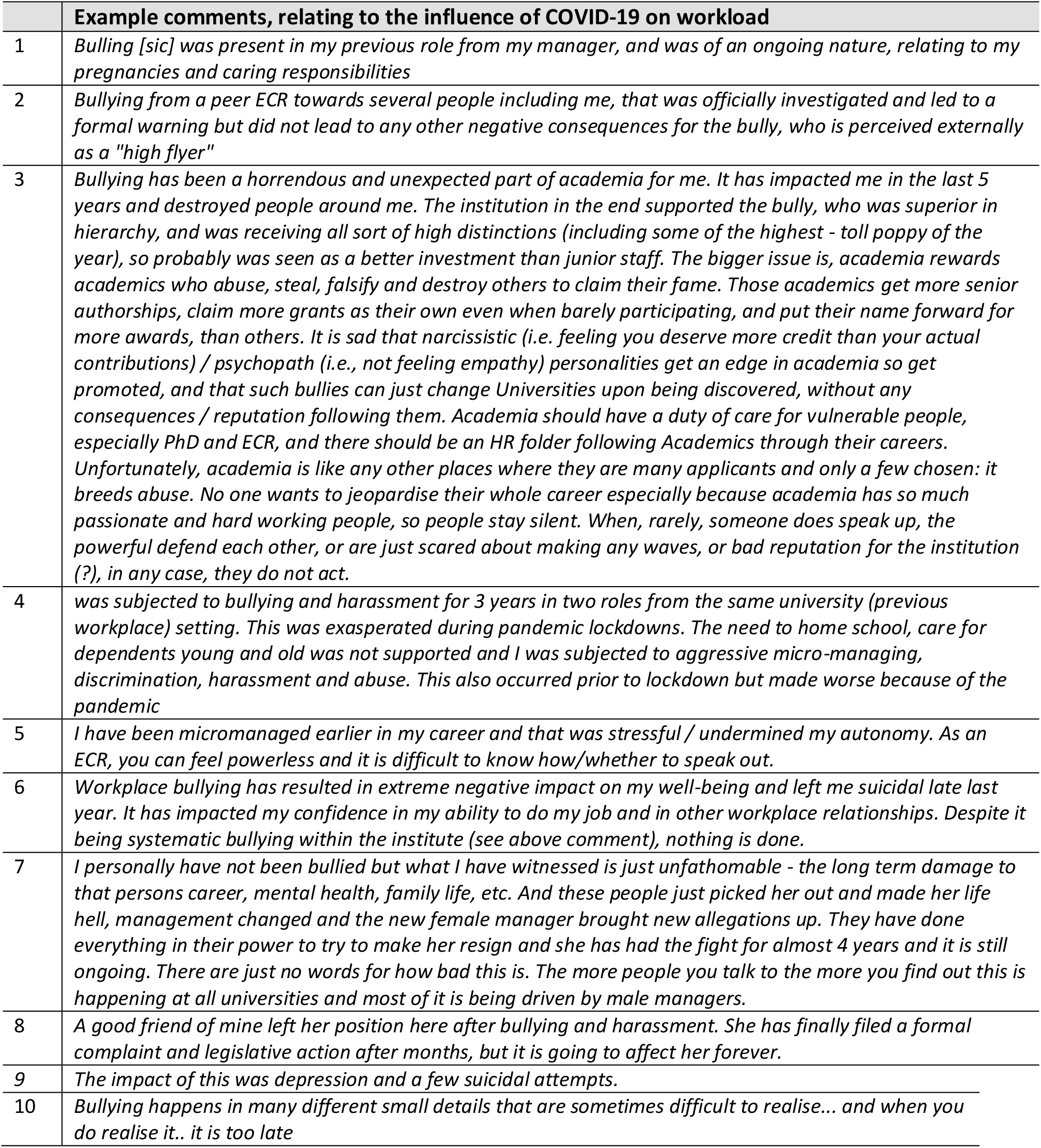
Comments from free text answers relating to bullying and harassment.

**Table 18.** Comments from free text answers relating to inequity and discrimination

|  | **Example comments, relating to inequity and discrimination** |
| --- | --- |
| 1 | *I have often been subjected to sexist comments, and doubts over the quality of my work because of my age and gender. Every woman in science I know has also.* |
| 2 | *I have also experience [sic] sexual harassment (unwanted attention and sexual propositions) and been stalked by the same student sporadically over the space of about 18 months.* |
| 3 | *I am a white, male early career researcher experiencing the pressure created by selection committees that intend to fill quotas rather than hiring talent. I feel especially bad for recent graduates who enter a deeply divisive job market in academia* |
| 4 | *Discrimination against young women on maternity leave- I have experienced it myself and have seen it happen to others. It's not even hidden! Plus the Prof asked a female researcher in an interview if she was going to have children, despite HR rep being in the room nothing was said. I was made casual and my income dropped (HR cc'd me in an email where it was discussed how to write my new position description to get away with this) after my first maternity leave. There are profs at the top bringing in grants who are untouchable and work by a different set of rules- no-one benefits from this scenario.* |
| 5 | *basically my 4 years PhD was a constant abuse and bullying by supervisor who hated working with " a man from the middle east". Stuff like "go back to your country". "you should be grateful you are here", "I gave you orders of what to do", "Men in the middle east dont [sic] respect women". The impact of this was depression and a few suicidal attempts. All of this was unprovoked. Misconduct, not by her, but another group who published research that was not reproducible by other members of the same lab* |
| 6 | *Every woman in science I know has also. I was doing a skilled job, and when I left I trained a less qualified male colleague to replace me. I later found out this less skilled and qualified man was earning 25% more than I was for a role I taught him how to do, when I had a PhD and he had an honours degree. I have so many of these stories, and I am just tired of fighting the same fights. I will be leaving academia at the first opportunity.* |
| 7 | *this is also an issue for those from cultures that do not value grandstanding. We can create opportunities for diversity, but there will always be closed doors so long as STEMM continues with a masculine euro-centric model of success.* |
| 8 | *As an Australian heterosexual male, I find myself subjected to discrimination within the research environment [sic]. Numerous training, career development, networking, and funding opportunities are being offered exclusively to females and other minority groups. I have been told that gender is considered in job applications submitted to my institution, with females preferentially selected for the purpose [sic] of creating a diverse workforce. Whether this is true or not, the current emphasis on females in STEMM has made me uncomfortable with my gender. I am tired of being told "you would have been perfect for that job/grant, if only you were female".* |
| 9 | *Women being blocked from promotion and career progression within the faculty* |
| 10 | *I think the research environment is highly selected towards masculine success. Women who succeed need to acquire masculine-type behaviours. Men are socialised towards competitiveness, assertiveness and occasional self-promotion whereas women are socialised towards teamwork, supportiveness and quiet achievement. I don't think any of the characteristics are wrong, but it does mean that most women must behave in a manner that feels unnatural/uncomfortable in order to be as visible in the workplace and have their work recognised.* |

### Questionable Research Practices

QRPs are also rife, 47% respondents reporting being impacted in 2022 compared with 38% in 2019. We asked respondents to indicate during what points in the research process they were pressured to engage in QRPs. Most common was with the ordering or inclusion of authors, with 49% of respondents claiming to be pressured to include or exclude authors (n = 443; see Supplementary Table 3 for more detail). Respondents reported pressure to consider engaging in QRPs during technical aspects of publications, including during Design/Methods (14.9%), during the analysis of data (17%), and during the presentation of results (18.4%). In each point in the publication process, greater than 5% of respondents were either uncertain or preferred not to say if they’d been pressured. Like other metrics, more women (54%) than men (40%) are subjected to pressure regarding authorship (*χ*^2^=11.24, df=3, P=0.011, no comparison with 2019 is available given we didn’t discriminate between different forms of QRPs in our earlier survey).

Next, we asked respondents more generally, if they were aware or suspicious of various forms of QRPs within their own Faculty (see Table 19). The most common practice reported was claiming of undeserved authorship (60.9%). Intertwined with this was the nearly equal exclusion of worthy co-authors (41.6%). Perhaps more damaging to the broad scientific enterprise is relatively high frequency that respondents reporting being aware of instances where fellow faculty fabricated data (made up data, 10%), plagiarized data (5.9%), falsified data (altered data, 8.4%), selectively dropped data sets from analysis without transparent explanation (26.5%), and trialed iterative statistical analysis until finding a model that yielded a “significant” result (45.5%). The cumulative outcome of these indiscretions no doubt is contributing to the so-called “reproducibility crisis” [8].

**Table 19.** Q46 – Awareness or suspicion of a QRPs by type within their faculty (n = 439-442)

| Question | Yes | No | Uncertain/<br>do not know | Do not want<br>to answer |
| --- | --- | --- | --- | --- |
| Fabricated data (made up data) | 10.0% | 71.2% | 18.4% | 0.5% |
| Plagiarized data | 5.9% | 73.0% | 20.6% | 0.5% |
| Falsified data (altered data) | 8.4% | 70.5% | 20.6% | 0.5% |
| Selectively dropped data from "outlier" cases without transparent explanation | 26.5% | 45.8% | 27.0% | 0.7% |
| Tried out a variety of different methods of analysis until one is found that yields a result that is statistically significant | 45.5% | 30.2% | 23.4% | 0.9% |
| Falsified biosketch, resume, reference list | 6.2% | 69.5% | 23.9% | 0.5% |
| Claimed undeserved authorship | 60.9% | 24.4% | 14.0% | 0.7% |
| Denied authorship to contributors | 41.6% | 41.4% | 16.1% | 0.9% |
| Used data without consent of other researchers | 20.2% | 60.5% | 18.6% | 0.7% |
| Been pressured by a study sponsor or contractor to engage in unethical research conduct or skewed presentation of research | 6.6% | 73.4% | 19.8% | 0.2% |
| Deliberately withheld data from the research community to gain personal or institutional advantage | 19.7% | 55.8% | 23.8% | 0.7% |
| Not disclosed a conflict of interest | 12.0% | 63.5% | 24.3% | 0.2% |
| Conducting research without appropriate ethical approval | 17.8% | 66.1% | 15.5% | 0.7% |

We asked respondents what they felt was the likely source of the pressure(s) leading to QRPs. The most common identified pressure point was “*my supervisor*” (24.6%), followed by “*colleagues in my faculty*” (15.1%), “*the competitive environment*” (10.2%), “*colleagues outside my faculty*” (7.5%), “*colleagues or managers at a former employer*” (4.4%), “*stakeholders with interest in the research*” (3.8%), “*a manager in my faculty*” (3.3%), or “the funder of the research” (3.2%). As with bullying and discrimination, the supervisor was most likely to be the perpetrator, followed by faculty colleagues. Again, supervisors and faculty cannot solve all problems, but they may be able to significantly influence the forces leading to QRPs.

Answers relating to respondent views about the severity of the QRPs and response from the institutions are shown in Table 20. It is concerning that 13.2% respondents felt that the nature of the QRPs of which they were aware are severe enough to warrant paper retraction, staff dismissal or a grant being repaid. A further 15.2% thought this was a possibility. At the same time, 20.1% believe there are on-going QRPs, commonly discussed by peers but are not being investigated. Only 7.9% felt that the described QRPs contributed “often” to the reproducibility crisis, while 47.6% of respondents felt that these QRPs “sometimes” contributed to the reproducibility crisis, which combined is more than half (55.5%). We consider these numbers in relation to the higher overall frequency of reported QRPs in Table 19; because disputed authorship is the most reported QRPs in Table 19, it is reasonable that respondents felt many of the QRP indiscretions would not necessarily compromise the validity of the reported data and therefore would be less likely to contribute to the reproducibly crisis. Table 21 includes statements from respondents describing QRPs at their own institution; these statements provide some context for the nature of QRPs and their severity.

**Table 20.** Questionable behaviours in the workplace (Questions 43, 44, and 45).

| Question | Yes | No | Maybe | Don't know |
| --- | --- | --- | --- | --- |
| If the nature of the questionable behaviour were known about by others, do you believe it would be viewed as sufficient to justify a paper retraction, dismissal or a grant being repaid? (n = 296) | 13.2% | 55.7% | 15.2% | 15.9% |
| Are you aware of on-going QRPs at your institute, or at a collaborating institute, that are commonly discussed within your peer group, but which you believe are not being investigated by institutional management? (n = 434) | 20.1% | 53.4% | 21.4% | 1.2% |
| How often do you believe you observe behaviour likely to contribute to the replication crisis at your institution? (n = 445) | **Often** | **Sometimes** | **Never** |  |
|  | 7.9% | 47.6% | 44.5% |  |

**Table 21.**
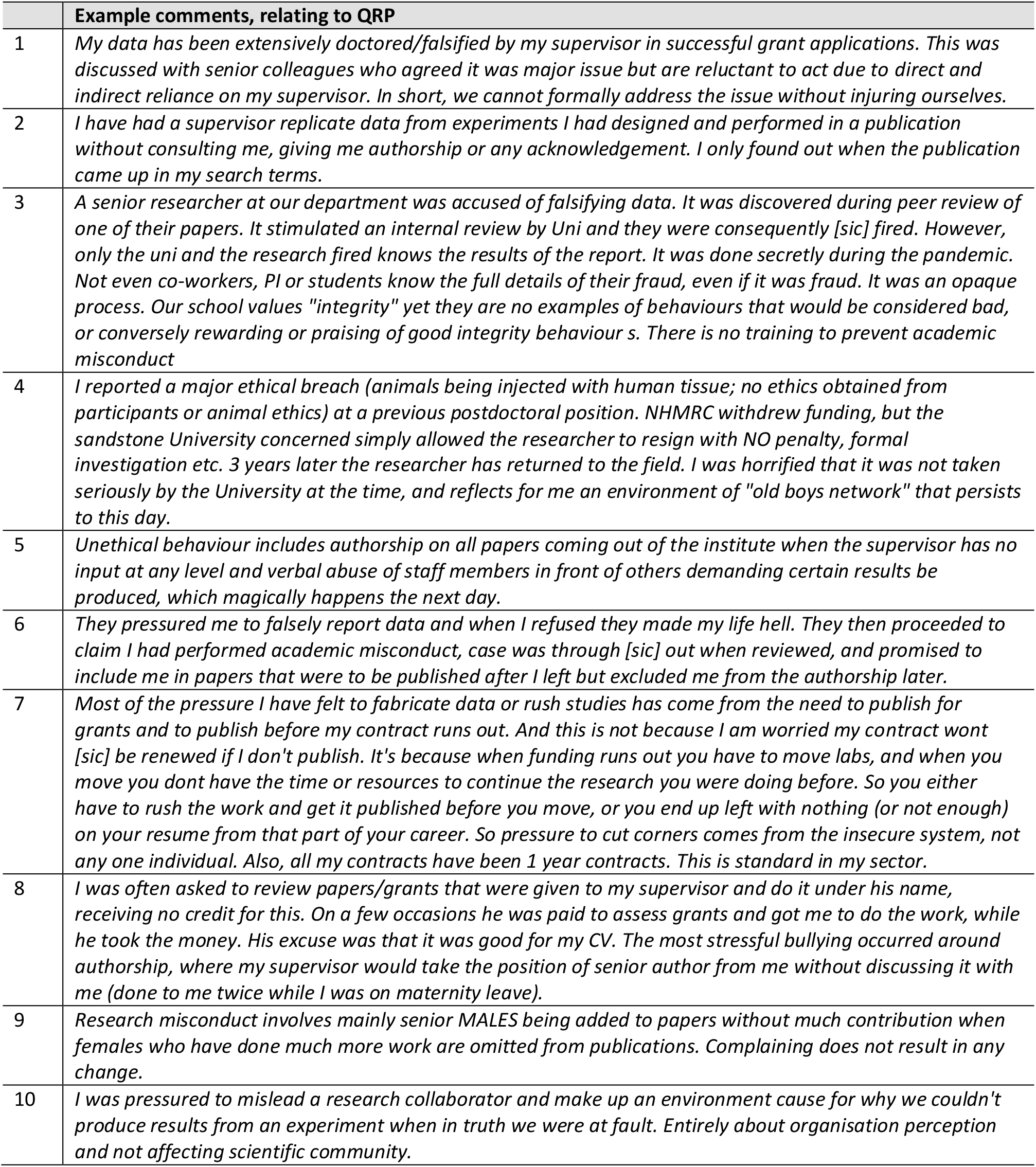
Comments from free text answers relating to questionable research practices

|  | Example comments, relating to QRP |
| --- | --- |
| 1 | *My data has been extensively doctored/falsified by my supervisor in successful grant applications. This was discussed with senior colleagues who agreed it was major issue but are reluctant to act due to direct and indirect reliance on my supervisor. In short, we cannot formally address the issue without injuring ourselves.* |
| 2 | *I have had a supervisor replicate data from experiments I had designed and performed in a publication without consulting me, giving me authorship or any acknowledgement. I only found out when the publication came up in my search terms.* |
| 3 | *A senior researcher at our department was accused of falsifying data. It was discovered during peer review of one of their papers. It stimulated an internal review by Uni and they were consequently [sic] fired. However, only the uni and the research fired knows the results of the report. It was done secretly during the pandemic. Not even co-workers, PI or students know the full details of their fraud, even if it was fraud. It was an opaque process. Our school values "integrity" yet they are no examples of behaviours that would be considered bad, or conversely rewarding or praising of good integrity behaviour s. There is no training to prevent academic misconduct* |
| 4 | *I reported a major ethical breach (animals being injected with human tissue; no ethics obtained from participants or animal ethics) at a previous postdoctoral position. NHMRC withdrew funding, but the sandstone University concerned simply allowed the researcher to resign with NO penalty, formal investigation etc. 3 years later the researcher has returned to the field. I was horrified that it was not taken seriously by the University at the time, and reflects for me an environment of "old boys network" that persists to this day.* |
| 5 | *Unethical behaviour includes authorship on all papers coming out of the institute when the supervisor has no input at any level and verbal abuse of staff members in front of others demanding certain results be produced, which magically happens the next day.* |
| 6 | *They pressured me to falsely report data and when I refused they made my life hell. They then proceeded to claim I had performed academic misconduct, case was through [sic] out when reviewed, and promised to include me in papers that were to be published after I left but excluded me from the authorship later.* |
| 7 | *Most of the pressure I have felt to fabricate data or rush studies has come from the need to publish for grants and to publish before my contract runs out. And this is not because I am worried my contract wont [sic] be renewed if I don't publish. It's because when funding runs out you have to move labs, and when you move you dont have the time or resources to continue the research you were doing before. So you either have to rush the work and get it published before you move, or you end up left with nothing (or not enough) on your resume from that part of your career. So pressure to cut corners comes from the insecure system, not any one individual. Also, all my contracts have been 1 year contracts. This is standard in my sector.* |
| 8 | *I was often asked to review papers/grants that were given to my supervisor and do it under his name, receiving no credit for this. On a few occasions he was paid to assess grants and got me to do the work, while he took the money. His excuse was that it was good for my CV. The most stressful bullying occurred around authorship, where my supervisor would take the position of senior author from me without discussing it with me (done to me twice while I was on maternity leave).* |
| 9 | *Research misconduct involves mainly senior MALES being added to papers without much contribution when females who have done much more work are omitted from publications. Complaining does not result in any change.* |
| 10 | *I was pressured to mislead a research collaborator and make up an environment cause for why we couldn't produce results from an experiment when in truth we were at fault. Entirely about organisation perception and not affecting scientific community.* |

Suspecting bullying and harassment and QRPs were likely to be found together in some workplaces, we examined that relationship. We found that for those who had experienced bullying, there was a higher incidence of impact by QRPs than for those who had not experienced bullying (authorship issues 57% versus 42%, presentation of results 26% versus 12%, analysis of data 25% versus 9% and Design/Method 22% versus 8%). Likewise, many more of those who had been impacted by QRPs had experienced bullying than those who had not (57% versus 34%). All these differences were statistically significant. (If have experienced bullying, differences with respect to Authorship, X^2^=11.33, df=3, P = 0.01; Design X^2^=21.38, df=3, P < 0.001; Analysis X^2^=24.19, df=3, P<0.001; Presentation X^2^=18.47, df=3, P < 0.001. If impacted by QRPs, differences with respect to bullying X^2^=18.34, df-2, P < 0.001.)

### Institutions do not act on complaints

Unfortunately, Australia does not have a centralized academic integrity office. Instead, institutions are required to investigate and manage such issues themselves. This introduces a conflict of interest where institutions, who generally want to avoid negative publicity, appear to be inclined to overlook or even cover up QRPs [26]. Thirty-three individuals remarked on inadequate institutional responses in the open text answers; these are illustrated by example comments provided in Table 22.

**Table 22:**
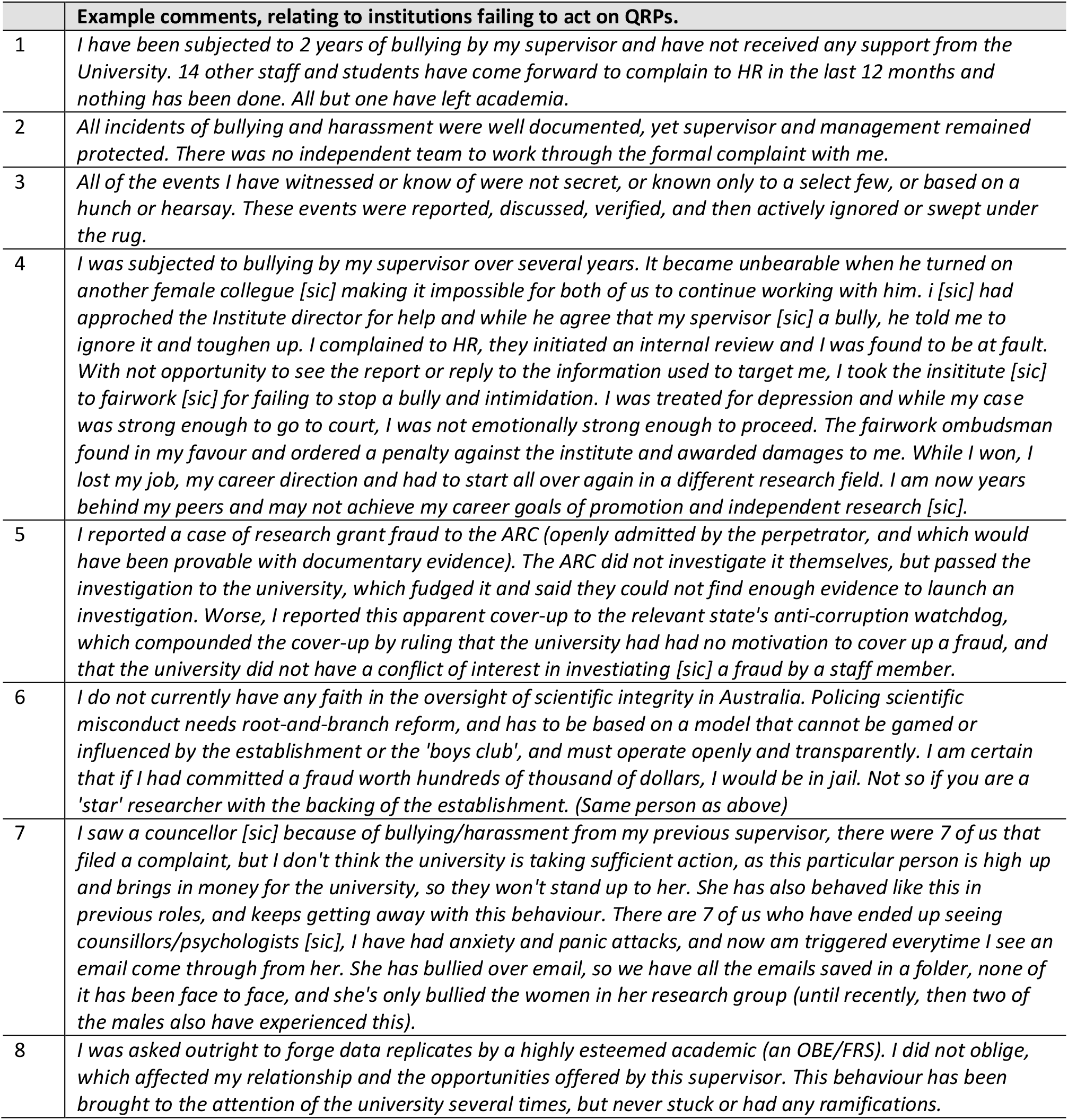
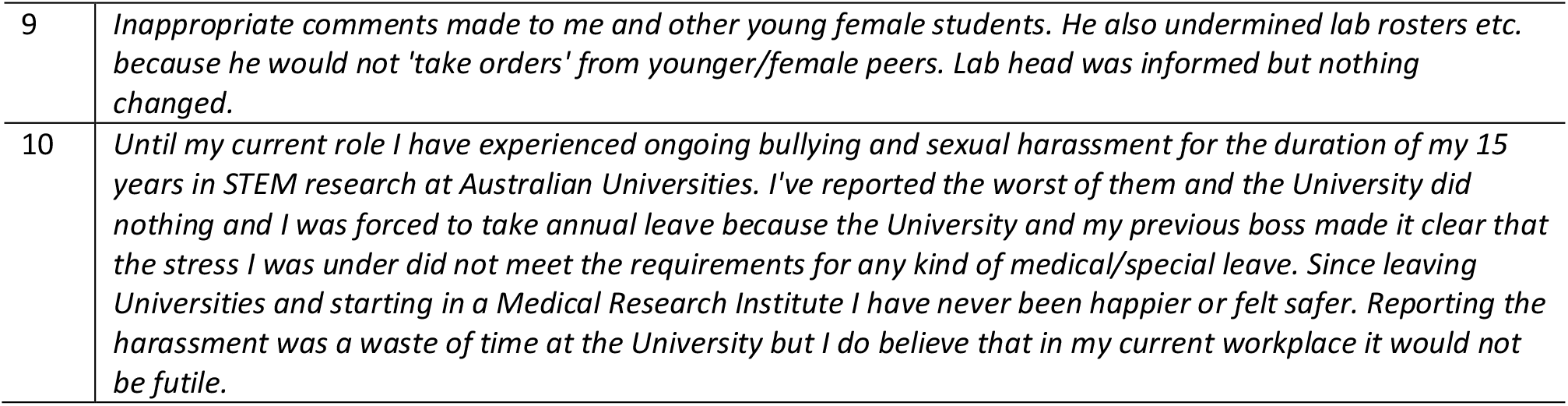
Comments from free text answers relating to Institutions failing to act on complaints.Discussion

|  | Example comments, relating to institutions failing to act on QRPs. |
| --- | --- |
| 1 | *I have been subjected to 2 years of bullying by my supervisor and have not received any support from the University. 14 other staff and students have come forward to complain to HR in the last 12 months and nothing has been done. All but one have left academia.* |
| 2 | *All incidents of bullying and harassment were well documented, yet supervisor and management remained protected. There was no independent team to work through the formal complaint with me.* |
| 3 | *All of the events I have witnessed or know of were not secret, or known only to a select few, or based on a hunch or hearsay. These events were reported, discussed, verified, and then actively ignored or swept under the rug.* |
| 4 | *I was subjected to bullying by my supervisor over several years. It became unbearable when he turned on another female colleague [sic] making it impossible for both of us to continue working with him. i [sic] had approached the Institute director for help and while he agree that my spervisor [sic] a bully, he told me to ignore it and toughen up. I complained to HR, they initiated an internal review and I was found to be at fault. With not opportunity to see the report or reply to the information used to target me, I took the insititute [sic] to fairwork [sic] for failing to stop a bully and intimidation. I was treated for depression and while my case was strong enough to go to court, I was not emotionally strong enough to proceed. The fairwork ombudsman found in my favour and ordered a penalty against the institute and awarded damages to me. While I won, I lost my job, my career direction and had to start all over again in a different research field. I am now years behind my peers and may not achieve my career goals of promotion and independent research [sic].* |
| 5 | *I reported a case of research grant fraud to the ARC (openly admitted by the perpetrator, and which would have been provable with documentary evidence). The ARC did not investigate it themselves, but passed the investigation to the university, which fudged it and said they could not find enough evidence to launch an investigation. Worse, I reported this apparent cover-up to the relevant state's anti-corruption watchdog, which compounded the cover-up by ruling that the university had had no motivation to cover up a fraud, and that the university did not have a conflict of interest in investiating [sic] a fraud by a staff member.* |
| 6 | *I do not currently have any faith in the oversight of scientific integrity in Australia. Policing scientific misconduct needs root-and-branch reform, and has to be based on a model that cannot be gamed or influenced by the establishment or the 'boys club', and must operate openly and transparently. I am certain that if I had committed a fraud worth hundreds of thousand of dollars, I would be in jail. Not so if you are a 'star' researcher with the backing of the establishment. (Same person as above)* |
| 7 | *I saw a counsellor [sic] because of bullying/harassment from my previous supervisor, there were 7 of us that filed a complaint, but I don't think the university is taking sufficient action, as this particular person is high up and brings in money for the university, so they won't stand up to her. She has also behaved like this in previous roles, and keeps getting away with this behaviour. There are 7 of us who have ended up seeing counsillors/psychologists [sic], I have had anxiety and panic attacks, and now am triggered everytime I see an email come through from her. She has bullied over email, so we have all the emails saved in a folder, none of it has been face to face, and she's only bullied the women in her research group (until recently, then two of the males also have experienced this).* |
| 8 | *I was asked outright to forge data replicates by a highly esteemed academic (an OBE/FRS). I did not oblige, which affected my relationship and the opportunities offered by this supervisor. This behaviour has been brought to the attention of the university several times, but never stuck or had any ramifications.* |
| 9 | *Inappropriate comments made to me and other young female students. He also undermined lab rosters etc. because he would not 'take orders' from younger/female peers. Lab head was informed but nothing changed.* |
| 10 | *Until my current role I have experienced ongoing bullying and sexual harassment for the duration of my 15 years in STEM research at Australian Universities. I've reported the worst of them and the University did nothing and I was forced to take annual leave because the University and my previous boss made it clear that the stress I was under did not meet the requirements for any kind of medical/special leave. Since leaving Universities and starting in a Medical Research Institute I have never been happier or felt safer. Reporting the harassment was a waste of time at the University but I do believe that in my current workplace it would not be futile.* |

## Discussion

World-wide reports of dissatisfaction with academic workplaces appear to be either growing or to be receiving greater visibility in the literature. In this report we aimed to dissect survey feedback solicited from Australian STEMM ECRs collected before the COVID-19 pandemic (2019) and after/during (2022). Data from the 2019 survey ECRs is published [1], and it identified that ‘love of science’ was a major career motivator for ECRs, but that job insecurity would likely force a career change.

The COVID-19 pandemic has exacerbated the stresses on Australian STEMM ECRs. These stresses further reveal systemic challenges within the academic ecosystem and emphasise the need for change. In 2019 we had 658 respondents [1] and in 2022 we had 530 respondents; the overall distribution of respondents within the various STEMM disciplines was similar. There were more female respondents (64%) than male (34%), and two most common age brackets were 31–35 years old (36.9%) and 36–40 years old (28.7%). While job satisfaction rates in the Australian workforce average 80% [16], ECR satisfaction in our surveys declined from 62% in 2019 to 57% in 2022. Almost half (48%) of respondents are stressed or very stressed daily, and many (58%) are considering leaving because of because of depression, anxiety, or other mental health concerns related to their work. Most (76%) agreed or strongly agreed that this is a poor time for a young person to commence a research career.

Academics are known to work long hours [18, 19]; a recent survey of academics in Australia and the UK found that respondents claimed to have worked a mean of 16–18 hours per week in excess of contract hours in the two weeks preceding the survey, and 90% reported working in excess of 10–12 hours per week over the previous six months [27]. Most of our respondents (61%) agreed in 2022 that their workload is too high compared with 49% in 2019. Of those who were employed full time and who worked at least 51 hours a week at work, 44% also work over 11 hours a week at home; 14% work more than 30 hours a week at home. These numbers correlated with many text responses that expressed concern over work-life balance. In inadequate job security as well as lack of funding or lack of independent positions were the dominant reasons cited for intending to leave research in 2019 (89.7 %) and 2022 (76.7 %). While the ecosystem became more competitive in 2022, an increasing number of respondents cited other reasons for potentially leaving, including stresses associated with poor work-life balance, demonstrating increasing load on the system.

It is estimated that because of the pandemic the university sector lost ~1.8 billion in revenue, and ~35,000 jobs (~25% academic positions and 75% administrative positions) [1, 9, 10]. While those in on-going positions are often sheltered from some of the stresses in academia, the pandemic created unusual circumstances and workloads where most felt vulnerable. Concerns over workload by staff were greatest for those on continuing positions (68.7%), then on short contracts <1 year (61.5%), and finally for those on 1-3 year contracts (56.4%). This relationship likely highlights the stress of short-term contracts, and perhaps workload being re-directed onto on-going staff when others did not have their contracts renewed. A Nature Careers article, *Pandemic burnout is rampant in academia*, cites an academic as not optimistic that workloads will ease any time soon, stating “*Every university will be under financial stringency, which means fewer faculty members and more workload*” [28].

In 2022, relative to 2019, ECRs reported increased rates of harassment from a someone in a position of power (33.5% increasing to 46.3%), lack of support from institutional superiors (60.0% to increasing to 70.5%) and being impacted by QRPs at their own institute (38.1% increasing to 47.4%). Bullying and harassment were reported by roughly half (47.8%) of respondents in 2022. The perpetrator was most frequently the supervisor (42.5%), another senior colleague (31.3%), or a peer (21.1%), and where motivation could be assigned, it was assumed to be related to the respondents’ gender (22.9%), race/ethnicity (9.7 %), age (11.0 %), nationality (7.7%), or class/socio-economic background (5.0%). We reason that high rates of bullying and harassment persist because almost half (44.3%) of ECRs do not believe that their institute would listen, and many (31.3 %) feel that they themselves would suffer negative consequences as an outcome of a complaint, compared to fewer (22.4%) who anticipated that their institute would act on a complaint.

The frequency that ECRs reported being impacted by QRPs at their own institute increased from 2019 (38%) to 2022 (47%). The most frequent (49%) complaints were associated with authorship, including both undeserved authorship (60.9%) and exclusion of worthy co-authors (41.6%). In the technical portion of manuscripts, ECRs reported pressure to questionably modify design/methods (14.9%), data analysis (17%), and presentation of results (18.4%). ECRs also indicated that they were aware of fellow faculty who had fabricated data (made up data, 10%), plagiarized data (5.9%), falsified data (altered data, 8.4%), selectively dropped data sets from analysis without transparent explanation (26.5%), and trialed iterative statistical analysis until finding a model that yielded a “significant” result (45.5%). It can be difficult to quantify an individual’s contribution to a specific research output, and we take the view that it is better to be inclusive rather than exclusionary in awarding authorship. From the COPE Authorship guidelines, “*Two minimum requirements define authorship across all definitions – making a substantial contribution to the work and being accountable for the work and its published form* [29]. The challenge with inappropriate authorship (inclusion or exclusion) is that publications are the primary currency of academia. This fact is also the motivator driving the high incidents of QRPs in the technical portions of publications, and in our view, this is the critical problem that underpins many of the challenges faced by ECRs in this study, and by academics around the world.

In academics’ efforts to create publication currency, a disconnect has formed where the true value or validity of a given publication has become difficult to quantify. The number of publications has grown to ~4.618 million in 2021 from 3.973 million in 2019, a 16% increase in only two years [30]. This surge in publication number potentially puts further downward pressure on the value of a publication, in a manner analogous to central banks printing money and putting downward pressure on the dollar. Regardless of the cause, it is critical to appreciate that high publication rates and concerns about declining publication quality pre-date the pandemic [31]. Concern over the validity or reliability of published data led to what has been termed *the reproducibility crisis* [8]. Some downplay the significance of the reproducibility crisis [32], but in 2011 Prinz *et al*., at Bayer, reported only being able to replicate 20-25% of 67 seminal studies [33]. They stated that despite well-resourced “*reasonable efforts (sometimes the equivalent of 3–4 full-time employees over 6–12 months), we have frequently been unable to reconfirm published data*.” A similar study was reported on in 2012 by Begley *et al*., from Amgen. Their team attempted to replicate 53 “*landmark*” cancer studies [34]. Despite efforts, including working with original authors to discuss discrepancies, exchange reagents or repeat experiments under the authors’ direction, only six (11%) could be reproduced. John Ioannidis who famously argued “*Why Most Published Research Findings Are False*” [35], critiqued 49 landmark medical publications from 1990-2003 with greater than 1,000 citations on the topics of hormone therapies, stents, aspirin and vitamin E. Of the 34 that had been replicated, 14 (41%) were incorrect or significantly exaggerated [36]. Fame, career stability and financial compensation are motivators for researchers to exaggerate claims in publications. For ECRs a single Cell, Nature or Science paper can change their career trajectory, potentially launching them ahead of those with a decade more experience; this is a potentially powerful and dangerous motivator that not surprisingly results in many high impact papers not being reproducible. Nations need become motivated to control and improve the reliability of the publication process. An economic analysis by Freedman and colleagues in 2015 estimated that non-reproducible pre-clinical research cost the USA $28 billion per year [37]; there is a risk that the growth in publication output may put downward pressure on actual meaningful scientific output, or at least obscure good science in a sea of non-reproducible publications.

ECRs are attempting to build a track record, and compete for grants and employment, also frequently through paper publication. In addition to the reproducibility flaws highlighted above, the field suffers broader challenges ranking ideas, and therefore researchers. In response to the challenge of assigning value to an output, in 2023, eLife, a prestigious journal, will no longer reject papers once they have entered the review process [38]; instead the paper, reviews, and responses to reviews will be published. Similarly, in response to the challenge of ranking the *innovative* aspect of grants [39], New Zealand is using a lottery system to award Explorer Grants [40]. In a survey of New Zealand researchers, most (63%) favored allocation of Explorer Grant funds via lottery, and interestingly many (40%) also favored application of this mechanism to other funding schemes [40]. Tangible and non-tangible qualities in a researcher are increasingly difficult to delineate; for example, it can be difficult to dissect quality from self-promotion as shown by a study of Academic Thoracic Surgeons in Canada and the USA that observed that Twitter activity was positively associated with a higher H-Index [41]. These data identified that “*the number of people followed (P = .048), and the frequency of tweeting (P = .046) as independent predictors of a higher h-index*”. This analysis does not demonstrate a causal relationship, nor that *Tweeting* confounds science, but it does suggest that a public profile may have a significant impact on the perceived value of published science, and thus the perceived capabilities of a given researcher. How many Twitter Followers read the papers published by these authors? What does a high Twitter follower count mean with respect to publication quality? We’re not sure, but we do intend to Tweet this paper once it has been published!

ECRs have also entered a minefield where prizes appear to play an increasing role in career success. There has been a corresponding boom in the number of prizes or awards in science [42], including many that target specific subgroups such as ECRs or women. While prestigious prizes, such as the Nobel Prize, are awarded based on a retrospective assessment of contributions, prizes given to PhD students or ECRs are often more prospective or maybe significantly influenced by their team or mentor’s contributions. An interesting analysis of tens of thousands of scientists found that specific mentorship was associated a 2-4-fold increase in a likelihood of prizewinning, but that later in their career protégés were most successful if their research evolved to differ from that of their mentor [43]. We wonder if awards or prizes might also distort metrics and put pressure on ECRs. In a recent editorial on this dilemma, Gabriel Popkin closed with the following comment “*One thing is certain: while scientists will keep winning prizes, what’s less clear is whether science itself is winning*.” [44]. Intertwined with prize winning and navigating the many “metrics challenges” of academic career development can be the growing perception that *luck* plays a critical role. A summary of interviews from those navigating academic careers repeatedly cited being *lucky* [45], while a similar summary of those navigating paths to become medical educators cited *serendipity* [46]. Similarly, there is growing data to suggest that the review processes used to rank grant applications are frequently underpowered, and winning can be assumed to be significantly influenced by chance or *luck* [39, 47]. In summary, today’s ECRs appear to be navigating a career path where success is intended to be based on merit, but where metrics are frequently distorted, and success can, in some cases, be attributed to luck.

The question is what to do about the impact of stresses on individual ECRs, on the field, and the possibility that stresses, like job security, may be directly impacting the validity of published literature? Based on our 2019 data [1], and these new data from 2022, we propose the following:

1. We need to develop an ecosystem that encourages institutions to invest in their ECRs, including their career development and stability. We need to slow the process down and take a long-term view of generating scientists and science. Standardised surveys could be used to collect data from PhD graduates and ECRs in Australia, identifying programs that have led to continuous employment, and institutions that supported ECR career development, making those institutions preferred places for study and employment. At the institutional level, group leaders could be encouraged to actively contribute to ECR development by integrating this output into workload calculations and promotion processes. Integrated into this ambition needs to be better alignment between Australia PhD graduate numbers and local job opportunities. Many graduates (75-78% in international surveys) aspire to obtain a job in academia [48, 49]. Unfortunately, in 2019, even before the pandemic, McCarthy and Wienk reported that since the mid-1990s Australian PhD graduate numbers have significantly outpaced academic jobs available in Australia [50]. Those who have an academic position, but who are dependent on grant funding to support their salary, are struggling, with low success rates for annual funding schemes (NHMRC Investigator Grant - 15.9% success rate in 2022, NHMRC IDEA Grant - 9.9% success rate in 2021, and the Australian Research Council Discovery Grant – 19 % success rate in 2022). Compounding poor academic employment prospects is the fact that Australian graduates more PhDs than the average of Organization for Economic Co-operation and Development (OECD) nations [51], yet has fewer advanced industries that typically employ highly skilled workers [52, 53]. One could argue that PhD students and ECRs are *cheap* labour for the research institutions, and that there are jobs overseas, but this does not seem like a sustainable lifestyle, nor a rational training investment for the country of Australia. It seems, instead, more rational to train a smaller number of stringently selected PhD students exceptionally well suiting them for both academic pursuits and alternate science career, thus offering these graduates more stable employment prospects, and more scientists for careers in Australia.
2. Aligned with the above goal of producing fewer PhD graduates, but with stronger skillsets, the PhD program in Australia could be extended from 3-years, to 4- or 5-years. It is increasingly recognized that graduates should have training appropriate to guide a future career in academia or industry [54], but this training takes time. In 2023, Australia will implement a new $206 million, 10-year, National Industry PhD Program that will link 1,800 PhD candidates with industry partners [55]. Students will be provided with 12 weeks of training to gain an understanding of industry and receive a stipend top-up for up to 4 years. The extended 4-year timeline is a movement in the right direction, but the graduate number should not be increased by an additional 1,800 graduates unless there is legitimate evidence of job market need. ECRs could also receive generic professional development training in skills which benefit in project management and leadership to better fit them for academia or the wider scientific workforce, as has been promoted in recent years by the Australian Academy of Science’s EMCR Forum [56].
3. Based on our data, which show growing concern over QRPs, an independent research integrity watchdog is needed in Australia. Professor David Vaux, an Australian immunologist and integrity expert, has argued that a national integrity watchdog is needed to remove the conflict-of-interest associated with institutions investigating themselves [57, 58]. Our respondents echo Vaux’s concerns, with only a fraction of ECRs (22.4%) indicating that they believed their institute would act on reported QRP concerns. An independent research integrity body could record and suggest general resolutions strategies for low level concerns, such as authorship disputes, and offer independent investigational resources for high level concerns, such as fraud [59, 60], which often comes hand in hand with bullying and harassment and has been linked to the volatile economic environment of the Australian higher education sector, and increased competition amongst its workforce [61]. In addition to higher quality investigation and being a better deterrent, this third-party watchdog could also better protect and anonymize whistle-blowers. University research and education is a billion-dollar business; it’s rational to assume there will be some bad behaviour and it’s rational to assume that a third-party oversight is needed. Finally, funding agencies should consider investing a portion of research funds into replication studies. This replication unit would randomly sample all publications funded by their agency, and these randomly selected publications should be scrutinized and replicated where possible. Perhaps just the possibility that publications might be replicated, would motivate researchers to take greater care in what data/claims were reported. While this process would consume some funds, and likely slow the rate of publication, it may increase the net value of published, thus adding value to the scientific endeavour.
4. Finally, it is time to be completely upfront with those considering entering PhD programs. Candidates should be made fully aware of discipline-specific career prospects. Institutions should be obligated to publish career and salary outcomes for previous graduates. There are plenty of reasons for a person to invest in educating themselves, but it is also a reasonable to expect institutions to provide the data required for potential candidates to conduct a cost-benefit analysis of such an investment.

## Conclusion

The pandemic has put immense stress on industries and professions, highlighting weaknesses and pushing many to the breaking point. Data from collected from ECRs working in STEMM disciplines across Australia suggest that the workplace culture that existed in 2019 [9] has further decayed over the pandemic. Job satisfaction has declined, workload concerns have increased, more ECRs have been impacted by QRPs, bullying and harassment, and in 2022 76% of ECRs believe “*now is a poor time to commence a research career*”. There are structural challenges in Australia, with PhD graduate numbers outstripping job opportunities as well as declining research funding, thereby contributing to career instability. As discussed above, structural stresses could be alleviated in part by transitioning away from a growth model that seeks to continuously generate more PhDs and more papers. At the institutional level, leaders may do more to protect their products (their scientists and their science) by slowing these processes down, and instead focusing on the provision stable work environments that aim to generate high quality reproducible outputs and which discourage poor behaviours or unethical practices. It’s now critical, for sake of the whole research ecosystem, that we reconsider how pressures may distort metrics and work together to identify new strategies to ensure that we promote the development of excellent scientists and excellent science.

## Methods

This research project explored challenges faced by ECRs in the sciences at universities and at independent research institutes in Australia; it was a follow up project conducted in 2022 to an earlier project conducted in 2019 [1].

The primary research questions from which the 2019 survey questions were derived were: (1) What are the principal factors that shape the ECR experience of various cohorts in the sciences in Australia? (3) What are the motivations for ECRs leaving the sciences? and (4) What are the specific features of the experiences and environment of those ECRs who remain in the sciences? For this follow up study, many of the questions were repeated and new questions were added to explore some aspects in greater depth. Emphasis was placed on the following questions: (1) What changes have been observed in the workplace culture and job satisfaction of ECRs in the sciences in Australia since the onset of the COVID-19 pandemic? (2) What is the extent, in the view of ECRs, of “bullying and harassment” and “questionable research practices” in their workplaces (3)? What is the nature, in the view of those ECRs, of “bullying and harassment” and “questionable research practices”; how do they impact them, and what is the response, if any, of the institutions. The definition of “early career researcher” for the purpose of this project included holding a PhD or equivalent, awarded no more than ten years prior and employment in an Australian university or independent research institute in a STEMM discipline.

### Ethics Approval

This study has been conducted according to the guidelines of the ethical review process of Queensland University of Technology (Approval Number 4846) and the National Health and Medical Research Council Statement on Ethical Conduct in Human Research.

### Survey

Survey questions are included in the Supplementary Data Section (Appendix 1). Quantitative data were collected from 530 eligible respondents in an on-line survey of ECRs working in a scientific environment in universities and research institutes across Australia. The questionnaire was developed by first compiling questions, often used in a broader or international context, from the 2019 on-line survey and from recent research literature, including questions from Wellcome Trust [11] and other recent surveys [62, 63] in order to cover all the themes covered by the research questions. Five additional questions were created, and validated, when no suitable question was identified elsewhere. These questions were combined to create a question bank of 64 questions for this survey relevant to the research questions and the Australian context, and the survey was pilot tested. Matters investigated include inequity, bias or discrimination with respect to age, gender, sexuality or race, bullying and harassment, questionable research practices, quality of supervision, career planning and professional development. The data from these questions were supplemented by questions seeking demographic information which included the gender, age, research discipline, country of origin, family situation and work arrangements of respondents.

The invitation to take part in the survey was distributed via email after direct contact with the institutions, via social media or “umbrella groups” such as EMCR Forum, Research Australia, The Australian Society for Medical Research (ASMR) with members or affiliates drawn from the STEMM community who were likely to include the target group.

A pilot study (n=16) permitted testing for understanding and clarity and to check for technical difficulties; The pilot survey ran from January 6 to January 11, 2022; no difficulties were found so the survey continued unchanged as the national survey. Data from this national survey is discussed in this paper. The national survey ran from January 12 to April 1, 2022. The survey was conducted online using Qualtrics XM. Eligibility to participate was determined by the initial questions in the survey.

### Data sharing

Full data sets will be shared upon request subject to the approval of the Queensland University of Technology Human Research Ethics Committee.

## Acknowledgements

We thank our survey respondents and all those who assisted in distributing invitations to the survey. MRD was supported by a National Health and Medical Research Council (NHMRC) of Australia Fellowship (APP1130013).

## Supplementary Tables

### Data for Figure 1

**Supplementary Table 1A:** Years post PhD.

| Answer (years) | 2019 | 2022 |
| --- | --- | --- |
| More than 10 years | 7.1 % | 0 % |
| 8-10 | 13.8 % | 22.8 % |
| 5-7 | 25.3 % | 31.9 % |
| 2-4 | 37.8 % | 36.49 % |
| 0-1 | 16 % | 8.9 % |

**Supplementary Table 1B:** Contract type.

| Answer | 2019<br>Raw count | 2019<br>Percent | 2022<br>Raw count | 2022<br>Percent |
| --- | --- | --- | --- | --- |
| University, teaching position | 20 | 3.1 % | 12 | 2.3 % |
| University, research only position and research institute | 399 | 62.5 % | 351 | 46.6 % |
| University, combined teaching and research position | 190 | 29.8 % | 134 | 25.3 % |
| University and hospital, combined clinical and research position | 29 | 4.5 % | 12 | 2.3 % |
| Other, please specify |  |  | 21 | 4 % |
| **Sum total** | **638** | **100 %** | **530** | **100 %** |

**Supplementary Table 1C:** Gender.S

| Answer | 2019<br>count | 2019<br>Percent | 2022<br>Count | 2022<br>Percent |
| --- | --- | --- | --- | --- |
| Man | 223 | 34.2 % | 182 | 34.3 % |
| Non-binary |  |  | 1 | 0.2 % |
| Woman | 430 | 65.8 % | 341 | 64.3 % |
| Prefer to self-describe |  |  | 0 | 0 % |
| Prefer not to say | 5 |  | 6 | 1.1 % |
| **Total** | **658** | **100 %** | **530** | **100 %** |

**Supplementary Table 1D:** Age.

| Answer | 2019<br>Raw count | 2019<br>Percent | 2022<br>Raw count | 2022<br>Percent |
| --- | --- | --- | --- | --- |
| **Less than 25** |  |  | 1 | 0.2 % |
| **25-30** | 109 | 16.5 % | 56 | 13.2 % |
| **31-35** | 282 | 42.7 % | 157 | 36.9 % |
| **36-40** | 171 | 25.9 % | 122 | 28.7 % |
| **40-45** | 42 | 6.4 % | 51 | 12 % |
| **Over 45** | 56 | 8.5 % | 38 | 8.9 % |
| **Total** | **660** | **100 %** | **425** | **100 %** |

**Supplementary Table 1E:** Where respondent was born.

| **Answer** | **2019**<br>**Raw count** | **2019**<br>**Percent** | **2022**<br>**Raw count** | **2022**<br>**Percent** |
| --- | --- | --- | --- | --- |
| **Australia** | 330 | 50.2 % | 208 | 48.9 % |
| **England** | 39 | 5.9 % | 25 | 5.9 % |
| **New Zealand** | 8 | 1.2 % | 12 | 2.8 % |
| **India** | 26 | 4 % | 11 | 2.6 % |
| **Italy** | 4 | 0.6 % | 6 | 1.4 % |
| **Vietnam** | 4 | 0.6 % | 3 | 0.7 % |
| **Philippines** | 3 | 0.5 % | 2 | 0.5 % |
| **China** | 18 | 2.7 % | 14 | 3.3 % |
| **Malaysia** | 11 | 1.7 % | 6 | 1.4 % |
| **Brazil** | 8 | 1.2 % | 8 | 1.9 % |
| **Other, please specify** | 207 | 31.5 % | 130 | 30.6 % |
| **Total** | 658 | 100 | 425 | 100% |

**Supplementary Table 1F:** Major countries within the “other” group

| **Answer** | **2019**<br>**Raw count** | **2019**<br>**Percent** | **2022**<br>**Raw count** | **2022**<br>**Percent** |
| --- | --- | --- | --- | --- |
| **USA** | 30 | 4.7 % | 15 | 3.5 % |
| **South Africa** | 8 | 0.5 % | 8 | 1.9 % |
| **Germany** | 18 | 2.7 % | 9 | 2.1 % |
| **France** | 11 | 1.7 % | 9 | 2.1 % |
| **Canada** | 10 | 1.5 % | 7 | 1.7 % |

**Supplementary Table 2.** Responses grouped by differences by country of birth

|  | 2019 |  |  | 2022 |  |  |
| --- | --- | --- | --- | --- | --- | --- |
|  | Born in Australia |  |  | Born in Australia |  |  |
|  | All<br>(n = 566,<br>647) | Yes<br>(n = 242,<br>316) | No<br>(n = 88,<br>109) | All<br>(n = 425) | Yes<br>(n = 208) | No<br>(n = 217) |
| Q13 Overall workload is too high | 48.6 % | 48.7 % | 47.7 % | 60.6 % | 59.6 % | 61.2 % |
| Q15 Overall satisfaction with job<br>(sat or v sat) | 62.3 % | 65.3 % | 55.7 % | 57.2 % | 62.0 % | 53.4 % |
| Q16-9 Do not feel safe | 12.5 % | 10.1 % | 13.6 % | 13.2 % | 12.5 % | 13.7 % |
| Q16-10 Satisfied with<br>commitment to diversity and<br>inclusiveness | 62.1 % | 69.6 % | 57.3 % | 56.4 % | 58.7 % | 54.8 % |
| Q19 Stressed or v stressed | NA |  |  | 48.4 % | 48.1 % | 48.6 % |
| Q22 Considered leaving for<br>mental health | NA |  |  | 57.5 % | 57.7 % | 57.3 % |
| Q27 Satisfied or v satisfied with<br>workplace culture | 51.0 % | 53.0 % | 44.7 % | 44.0 % | 49.0 % | 39.9 % |
| Q28 Poor time for a young person<br>to begin – agree & strongly agree | 64.7 % | 66.1 % | 67.0 % | 76.0 % | 77.4 % | 74.9 % |
| Q30-1 inequitable hiring practices | 38.5 % | 37.3 % | 47.4 % | 49.3 % | 40.9 % | 55.9 % |
| Q30-2 Harassment based on<br>power position | 33.5 % | 37.2 % | 36.1 % | 46.3 % | 43.8 % | 48.3 % |
| Q30-3 Lack of support from<br>institutional superiors | 60.0 % | 63.5 % | 60.9 % | 70.5 % | 64.4 % | 75.3 % |
| Q30-4 Impacted by QRPs in<br>institution sometimes & often | 38.1 % | 36.2 % | 46.4 % | 47.4 % | 46.2 % | 48.3 % |

**Supplementary Table 3.** Q41 - Exposure to unethical pressure by nature of QRPs (n = 433).

| Question | Yes | No | Uncertain/do<br>not know | Prefer not to<br>answer |
| --- | --- | --- | --- | --- |
| Ordering/inclusion of authors | 49.0% | 41.8% | 7.2% | 2.0% |
| Design/method | 14.9% | 78.7% | 5.0% | 1.4% |
| Analysis of data | 17.0% | 77.2% | 4.8% | 1.1% |
| Presentation of results | 18.4% | 76.0% | 4.5% | 1.1% |

## Appendix 1 National On-line Survey Questions

Source of questions, if taken from elsewhere, appears in brackets. Questions without source are the same as the previous survey

### Eligibility

1. Which of the following best describes your current position within the STEMM <u>research community</u>? *By research community, we are referring to <u>all those who conduct or support research</u>*. (Wellcome Trust, 2020)

i. I am a student –[Terminate these]
ii. I am employed / contracted / freelance
iii. I am taking a career-break / on leave (e.g. parental)
iv. I am looking for work / unemployed
v. I am retired
vi. I used to be part of the research community, but no longer am
vii. I have <u>never</u> been part of the research community–[Terminate these]
viii. Other, please specify
2. Do you have a PhD or doctoral qualification?

i. Yes
ii. No-Terminate these
iii. Currently studying towards this level of qualification –[Terminate these]
3. What is the number of years since completion of your highest degree?

i. 0–1
ii. 2–4
iii. 5–7
iv. 8–10
v. More than 10 years – [terminate these]
4. What is the nature of your employment?

i. University, teaching position
ii. University, research only position
iii. University, combined teaching and research position
iv. University and hospital, combined clinical and research position
v. Government research institute (e.g. CSIRO, ANSTO) - [terminate these]
vi. Research institute
vii. Not for profit organisation – [terminate these]
viii. Other, please specify

### Demographics

5. Which of the following best describes your gender? **(Wellcome Q58)**

i. Man
ii. Non-binary
iii. Woman
iv. Prefer to self-describe
v. Prefer not to say
6. What is your primary research discipline? Select the appropriate Australian FOR code:

i. Division 01 Mathematical Sciences
ii. Division 02 Physical Sciences
iii. Division 03 Chemical Sciences
iv. Division 04 Earth Sciences
v. Division 05 Environmental Sciences
vi. Division 06 Biological Sciences
vii. Division 07 Agricultural and Veterinary Sciences
viii. Division 08 Information and Computing Sciences
ix. Division 09 Engineering
x. Division 10 Technology
xi. Division 11 Medical and Health Sciences

If answered **1iv)**

Why did you leave the research community?

i. I’m no longer interested in a research-related career
ii. I wanted to apply my skills elsewhere
iii. My contract ended / my role was terminated
iv. Too difficult to find a job / insecure career path
v. Too difficult to obtain funding
vi. The career was too demanding
vii. For career progression / development
viii. For better compensation / salary
ix. For a better work-life balance
x. It was impacting on my wellbeing and mental health
xi. To launch my own business
xii. Personal reasons
xiii. Retirement
xiv. Bullying and harassment
xv. Discrimination
xvi. Other, please specify
xvii. I’d prefer not to say

Then terminate

### About your job and work status and workload

7. In which manner are you employed:

i. Full time continuing
ii. Part time continuing
iii. Full time fixed term contract
iv. Part time fixed term contract
v. Contractor / self employed
vi. Other (please specify)
8. If you are on a fixed term contract, what is the total length of your [fixed-term] contract?

i. Less than 1 year
ii. 1 to three years
iii. More than 3 years (please specify in comment)
iv. Other (please specify in comment)
9. What is your employment fraction? (i.e. 0.2 =one day per week)

i. 0.2 FTE
ii. 0.4 FTE
iii. 0.5 FTE
iv. 0.6 FTE
v. 0.8 FTE
vi. 1.0FTE
vii. Other, please explain (please specify in comment)
10. On average, how many hours per week do you work in your workplace, including in field or clinical settings?

i. Up to 20
ii. 21-30
iii. 31-40
iv. 41-50
v. 51-60
vi. 61-70
vii. Greater than 70
11. On average, how many hours per week do you undertake work related to your employment at home?

i. Up to 5 hours
ii. 6-10 hours
iii. 11-15 hours
iv. 16-20 hours
v. 21-30 hours
vi. Greater than 30 hours
vii. Other, please specify (please specify in comment)
12. Has this time working at home changed due to COVID-19? (new question)

i. Yes, I now spend more time working at home
ii. Yes, I now spend less time working at home
iii. No
13. How would you describe your overall workload? much too low, about right, too high A postdoctoral appointment, or “postdoc,” is a temporary position awarded in academe, industry, government or a non-profit organization primarily for gaining additional education and training in research. For the next question, please include any position you consider to be a “postdoc” even if your employer did not or does not. Please also count reappointments to the same position as one appointment.
14. How many postdoctoral appointments have you had, including your current position if applicable? Select one. If “other” please explain.

i. 1
ii. 2
iii. 3
iv. More than 3 (please specify in Other)
v. Other (please specify in comment)

### Job satisfaction

15. How would you rate your overall satisfaction with your current job? 5 point scale very satisfied to very dissatisfied
16. To what extent do you agree with the follow statements about your current job?

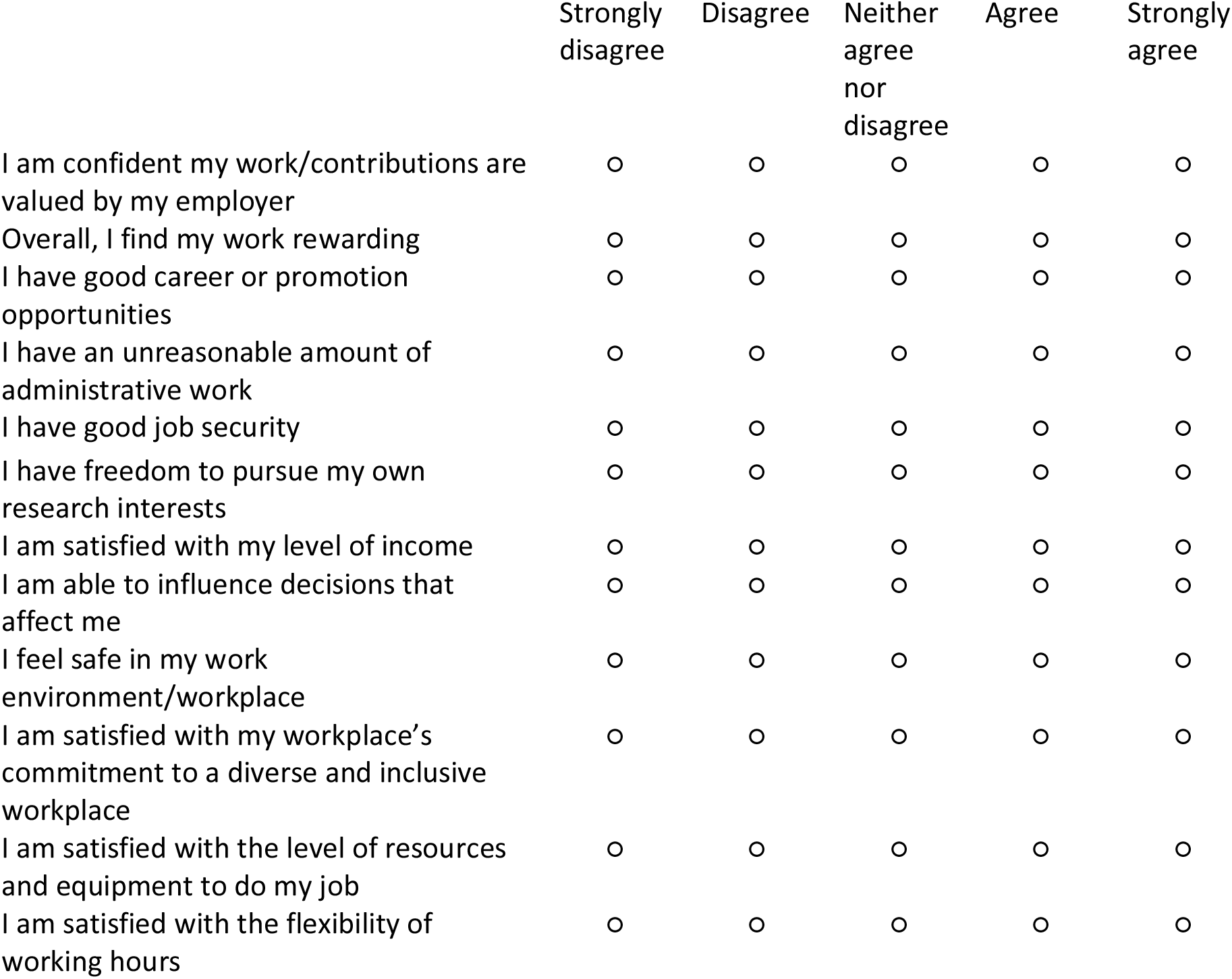
17. If there was one factor you could change that would make a major difference to your levels of job satisfaction what would it be? (Select one ONLY)

|  | Strongly<br>disagree | Disagree | Neither<br>agree<br>nor<br>disagree | Agree | Strongly<br>agree |
| --- | --- | --- | --- | --- | --- |
| I am confident my work/contributions are valued by my employer | ☐ | ☐ | ☐ | ☐ | ☐ |
| Overall, I find my work rewarding | ☐ | ☐ | ☐ | ☐ | ☐ |
| I have good career or promotion opportunities | ☐ | ☐ | ☐ | ☐ | ☐ |
| I have an unreasonable amount of administrative work | ☐ | ☐ | ☐ | ☐ | ☐ |
| I have good job security | ☐ | ☐ | ☐ | ☐ | ☐ |
| I have freedom to pursue my own research interests | ☐ | ☐ | ☐ | ☐ | ☐ |
| I am satisfied with my level of income | ☐ | ☐ | ☐ | ☐ | ☐ |
| I am able to influence decisions that affect me | ☐ | ☐ | ☐ | ☐ | ☐ |
| I feel safe in my work environment/workplace | ☐ | ☐ | ☐ | ☐ | ☐ |
| I am satisfied with my workplace's commitment to a diverse and inclusive workplace | ☐ | ☐ | ☐ | ☐ | ☐ |
| I am satisfied with the level of resources and equipment to do my job | ☐ | ☐ | ☐ | ☐ | ☐ |
| I am satisfied with the flexibility of working hours | ☐ | ☐ | ☐ | ☐ | ☐ |

o Improved working hours
o More protected time for research
o Improved leave provisions
o Improved institutional / organisational culture
o Improved promotional opportunities
o Better pay
o Improved job security
o Improved mentorship / supervision
o More family friendly environment
o Support for career development
o Other (please specify)
o None of these. I am very satisfied with my current job

### About personal challenges which relate to your work

18. Do you have any caring responsibilities? (**Wellcome Q63)** Select all that apply

i. None
ii. Primary carer of a child/children (under 18)
iii. Primary carer of disabled child/children
iv. Primary carer or assistant for a disabled adult (18 years or over)
v. Primary carer or assistant for an older person/people (65 and over)
vi. Secondary carer (another person carries out the main caring role)
vii. Prefer not to say
19. How stressed do you feel at work / while working on an average day? (**Wellcome)** Grid question 5-point scale 1 = Not at all stressed, 4= Neutral, 7 = Extremely
**20.** Do you have any of the following disabilities, long-term health conditions, mental health conditions or impairments? *Please select all that apply*. **(Wellcome Q67)**

i. No known disability, long-term condition or impairment
ii. Dyslexia
iii. Other neurodiverse (such as dyscalculia, autism)
iv. Hearing
v. Speech
vi. Visual
vii. Long-term health condition (such as diabetes, Multiple Sclerosis, heart condition, epilepsy, energy-limiting conditions, chronic pain)
viii. Mental health
ix. Mobility
x. Musculoskeletal (including back, neck and shoulder)
xi. Listed above but prefer not to specify
xii. Other, please specify
xiii. Prefer not to say
21. Do you experience barriers or limitations in your day-to-day activities related to any of your health conditions, impairments or disabilities? **(Wellcome Q68)**

i. Not applicable
ii. No
iii. Yes
iv. Prefer not to say
22. Have you considered leaving science because of depression, anxiety, or other mental health concerns related to your work? **Nature postdoc satisfaction survey 2020** (Nature Research & Shift Learning, 2020)

i. Yes
ii. No
iii. I’d prefer not to say
23. Do you face any barriers in achieving a successful career in the research community? *Please select all that apply. (****W*ellcome)**

i. Lack of funding
ii. Lack of training in relevant skills
iii. Lack of training in relevant field
iv. Unmanageable workload
v. Lack of advice and guidance
vi. Lack of support from institution/workplace
vii. Job insecurity
viii. Bullying and harassment
ix. Lack of opportunities
x. Inability to relocate
xi. Caring responsibilities
xii. Inequalities / discrimination / bias
xiii. None of the above
xiv. Other, please specify
24. Did your role change significantly as a result of COVID-19 during 2020 or 2021? Please select all that apply. (Schultz, 2020 with 2021 added)

i. reduced hours (e.g. full time to part time)
ii. became unemployed
iii. changed proportion of research/teaching
iv. unable to supervise research projects
v. other
25. How stable was the STEMM workforce before the COVID-19 pandemic in your professional area? Please rate on a scale of 1 to 10 unstable to highly stable (adapted from AustAssnScience)
26. How stable is the STEMM workforce now since the COVID-19 pandemic in your professional area? Please rate on a scale of 1 to 10 unstable to highly stable (new question)

### About overall research culture

27. How satisfied are you with the culture of your workplace very satisfied to very dissatisfied
28. How do these statements following correspond with your views about the nature of your job? 5 point scale rating from strongly agree to strongly disagree for each:

i. This is a poor time for any young person to begin an academic career in my field.
ii. If I had it to do over again, I would not become an academic
iii. My job is a source of considerable personal strain
29. To what extent have the following characteristics of your workplace culture impacted you or your career advancement? VERY SUPPORTIVE-SUPPORTIVE - NEITHER SUPPORTIVE NOR A PROBLEM – NOT SUPPORTIVE/A PROBLEM – VERY UNSUPPORTIVE/ A MAJOR PROBLEM - NOT APPLICABLE

i. Level of support from supervisor/manager in applying for promotion
ii. Guidance received in performance reviews
iii. Opportunities for professional development
iv. Opportunities to undertake/complete qualifications
v. Access to research funding
vi. The attitude towards people of my age
vii. The attitude towards people of my gender
viii. The attitude towards people of my ethnic background
ix. The attitude towards people of my sexual orientation
x. Availability of informal mentoring
30. To what extent have the following negative characteristics of some workplace cultures impacted you or your career advancement in your workplace? NEVER A PROBLEM - SOMETIMES A PROBLEM - A SIGNIFICANT PROBLEM

i. Inequitable hiring practices
ii. Harassment based on different power position
iii. Lack of support from institutional superiors
iv. Questionable research practices of colleagues within my institution
v. Questionable research practices of colleagues outside my institution
31. Thinking about the last job you left, what was the reason for leaving? (tick all that apply)

i. Lack of funding for new contract/further employment
ii. Career progression / development
iii. The new job is better suited to my interests / skills
iv. For better compensation / salary
v. or full-time permanent position
vi. Better work-life balance
vii. Unhappy with role
viii. Looking to relocate / partner was relocated
ix. Launch my own business
x. Terminated / made redundant
xi. Maternity / paternity leave
xii. Retired
xiii. Personal reasons
xiv. Unhappy with organisational culture
xv. I was subjected to bullying or harassment at work
xvi. I’d prefer not to say
xvii. Not applicable
xviii. Other, please specify
32. How far do you agree or disagree with the following statements relating to your current working environment? **(Wellcome)** Grid question 5-point scale

i. My working environment promotes a good work-life balance
ii. My working environment promotes a collaborative culture
iii. Creativity is welcomed within my working environment in all its forms
iv. Healthy competition is encouraged within my working environment
v. Unhealthy competition is present within my working environment
vi. My institution/workplace values speed of results over quality
vii. My institution/workplace could do more to ensure research practices do not cut corners
viii. Rigour of results is considered an important research outcome by my institution/workplace
ix. My institution/workplace places more value on meeting metrics, than it does on research quality
x. I am confident that my institution/workplace would listen and take action if I raised a concern
xi. The culture around research in my working environment supports my ability to do good quality research
xii. My institution/workplace’s expectations of me to undertake a number of roles leaves me little time for research
xiii. My working environment hinders researchers getting on with their research
xiv. My institution/workplace provides me with support to navigate the grant application process

### About Bullying and Harassment

33. During your research career have you ever…? (**Wellcom**e) Yes, No, Prefer not to say, N/A

i. Experienced bullying or harassment
ii. Witnessed bullying or harassment
**34.** If you have <u>experienced</u> bullying or harassment, who was the perpetrator(s)? (**Wellcome)** Select all

i. Supervisor or manager
ii. Other senior colleague
iii. A peer
iv. Other, please specify
v. Prefer not to say
**35.** If you have <u>witnessed</u> bullying or harassment, who was the perpetrator(s)? (**Wellcome)** Select all

i. Supervisor or manager
ii. Other senior colleague
iii. A peer
iv. Other, please specify
v. Prefer not to say
36. During your research career have you ever… (**Wellcome)** Yes, No, Prefer not to say, N/A

i. Experienced discrimination
ii. Witnessed discrimination
37. In cases where you have <u>experienced</u> bullying and harassment or discrimination, was this behaviour related to… (**Wellcome)** Select all

i. Age
ii. Class / socio-economic background
iii. Disability
iv. Gender
v. Gender identity (e.g. trans or non-binary)
vi. Nationality
vii. Race or ethnicity
viii. Religion
ix. Sexual orientation
x. Other, please specify
xi. Prefer not to say
xii. N/A
38. In cases where you have <u>witnessed</u> bullying and harassment or discrimination, was this behaviour related to… (**Wellcome**) Select all

i. Age
ii. Class / socio-economic background
iii. Disability
iv. Gender
v. Gender identity (e.g. trans or non-binary)
vi. Nationality
vii. Race or ethnicity
viii. Religion
ix. Sexual orientation
x. Other, please specify
xi. Prefer not to say
xii. N/A
39. Within your workplace, do you feel your concerns relating to experiences of bullying and/or discrimination would be…? (**Wellcome)** Yes, No, Unsure

i. Listened to
ii. Appropriately acted upon
40. Would you feel comfortable speaking out about instances of bullying and/or discrimination without negative personal consequences from within your workplace? (**Wellcome)**

i. Yes
ii. No
iii. Unsure
iv. Prefer not to say

### About prevalence and impact of research misconduct

41. Have you ever been exposed to unethical pressure concerning (**Horbac**h adapted) Yes No Uncertain/do not know Do not want to answer Not applicable for each

i. Ordering/inclusion of authors
ii. Design/method
iii. Analysis of data
iv. Presentation of results
**42.** If you answered ‘yes’ to having been exposed to unethical pressure; please indicate the sources of the pressure (choose all that apply): (**Horbach adapted)**

i. The funder of the research
ii. Stakeholders with interest in the research
iii. My supervisor
iv. Colleagues in my faculty
v. A manager in my faculty
vi. Colleagues outside my faculty
vii. Colleagues or managers at a former employer
viii. The competitive environment
ix. Not applicable
x. Other, please specify If you wish to describe any instance of research misconduct which has impacted you may do so at the end of the survey
43. If the nature of the questionable behaviour were known about by others, do you believe it would be viewed as sufficient to justify a paper retraction, dismissal or a grant being repaid? (new question) Yes no maybe don’t know
44. Are you aware of on-going questionable research practices at your institute, or at a collaborating institute, that are commonly discussed within your peer group, but which you believe are not being investigated by institutional management? Yes no Don’t know Prefer not to say Other
45. The “replication crisis” is a now well described phenomenon used to describe situations where it has not been possible to replicate published studies (Ioannidis, 2005). How often do you believe you observe behaviour likely to contribute to the replication crisis at your institution? Never Sometimes Often
46. Have you known about or justifiably suspected that any of the colleagues in your faculty during your time as a researcher has (Horbach et al., 2020 adapted) Yes No Uncertain/do not know Do not want to answer Not applicable

i. Fabricated data
ii. Plagiarized data
iii. Falsified data
iv. Selectively dropped data from “outlier” cases without transparent explanation
v. Tried out a variety of different methods of analysis until one is found that yields a result that is statistically significant
vi. Deliberately withheld data from the research community to gain personal or institutional advantage
vii. Falsifed biosketch, resume, reference list
viii. Not disclosed a conflict of interest
ix. Claimed undeserved authorship
x. Denied authorship to contributors
xi. Been pressured by a study sponsor or contractor to engage in unethical research conduct or skewed presentation of research
xii. Used data without consent of other researchers
xiii. Conducting research without appropriate ethical approval

### About Leadership

47. How satisfied are you with the leadership and management of your workplace? Very satisfied to very dissatisfied
48. To what extent do you agree or disagree with the following statements regarding your institutional senior management? **(Wellcome)** Grid question 5-point scale

i. I think senior management makes wise decisions
ii. I am satisfied with the way my institution/workplace handles performance reviews
iii. Leaders communicate clear expectations regarding behaviours and/or culture in my working environment
49. How successful is your <u>workplace team</u> in demonstrating each leadership characteristic? **(Wellcome)** Grid question Extremely unsuccessful, Somewhat unsuccessful, Neutral, Somewhat successful, Extremely successful, I don’t know, N/A

i. Setting the direction for research and creating the plans and systems to achieve it
ii. Leading and supporting teams of diverse individuals
iii. Setting and upholding standards in the conduct of research and its application
iv. Creating development and career opportunities
v. How successful is your institution / workplace as a whole in demonstrating each leadership characteristic? **(Wellcome)** Grid question Extremely unsuccessful, Somewhat unsuccessful, Neutral, Somewhat successful, Extremely successful, I don’t know, N/A

i. Setting the direction for research and creating the plans and systems to achieve it
ii. Leading and supporting teams of diverse individuals
iii. Setting and upholding standards in the conduct of research and its application
iv. Creating development and career opportunities

### About supervision and mentoring

**51.** How satisfied are you with the support for your career development/professional development? Very satisfied to very dissatisfied
52. When you started with your current employer how useful did you find the following?

53. Has your supervisor, PI or manager done any of the following within the last 12 months? *Please select all that apply*. **(Wellcome)**

|  | Not at<br>all useful | Not very<br>useful | Neither<br>useful or<br>not | Useful | Extremely<br>useful | Offered<br>but not<br>taken | Not<br>offered |
| --- | --- | --- | --- | --- | --- | --- | --- |
| Institutional-wide<br>induction<br>programs | ☐ | ☐ | ☐ | ☐ | ☐ | ☐ | ☐ |
| Departmental<br>/Faculty/Unit<br>induction<br>program | ☐ | ☐ | ☐ | ☐ | ☐ | ☐ | ☐ |
| The local<br>induction to your<br>current role | ☐ | ☐ | ☐ | ☐ | ☐ | ☐ | ☐ |

i. Had a conversation with you about your career aspirations
ii. Provided career advice and guidance
iii. Discussed your performance
iv. Provided an example of appropriate ethical codes
v. Noted your achievements
vi. Offered you training to support your skill development
vii. Provided an example of appropriate research standards
viii. Connected you to others within or outside your field
ix. Supported you with personal issues
x. Supported your wellbeing
xi. Provided expert advice
xii. Conducted a formal appraisal
xiii. Discussed alternative career options
xiv. Requested your feedback on their management of you
xv. None of the above
xvi. Not applicable
54. To what extent do you agree or disagree with the following statements regarding the management of your work? **(Wellcome)** Grid question 5-point scale 1=Strongly disagree, 4= Neither disagree nor agree, 7= Strongly agree, Not applicable

i. My supervisor regularly reviews my work
ii. I would feel comfortable approaching my supervisor if I couldn’t reproduce lab results
iii. My supervisor values negative results that don’t meet an expected hypothesis
iv. I have felt pressured by my supervisor to produce a particular result
v. My supervisor gives me freedom to explore my results
**55.** On average, how much one-on-one contact time do you spend with your supervisor/PI each week? (**Wellcome)**

i. Less than an hour
ii. Between one and three hours
iii. More than three hours
iv. Other, please specify
v. N/A
56. Have you participated in your institution’s staff review/appraisal scheme in the last two years? How would you rate this scheme’s usefulness? Not at all useful to extremely usedful, NA

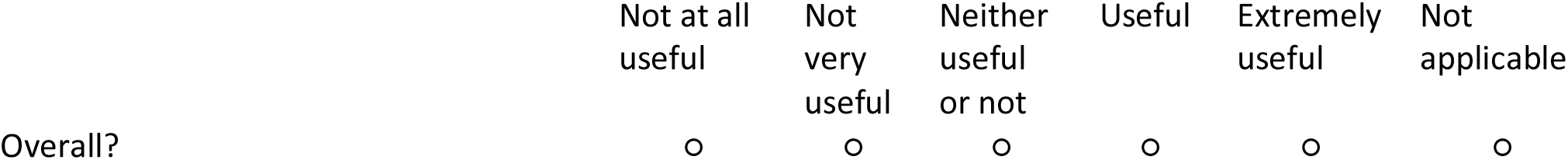

### About whether you are considering a change in your work or career

57. What would be the main reason you would consider leaving a career in research? If “Other” please specify

i. Family/carer responsibilities
ii. Interpersonal problems with your supervisor
iii. Inadequate job security
iv. A lack of independent positions available
v. A lack of funding
vi. Other (please specify)
58. Does your institute have career advisory services for science ECRs?

i. Yes, but I haven’t had any contact with them
ii. Yes, and their offerings have been useful
iii. Yes, but their offerings have not been useful
iv. No
v. I don’t know

### A little more about you

59. Which of the following best describes your sexual orientation? **(Wellcome Q60.)**

i. Asexual
ii. Bi/bisexual
iii. Gay man
iv. Gay woman/lesbian
v. Heterosexual/straight
vi. Queer
vii. Prefer not to say
viii. Other, please specify
60. What is your age?

i. Less than 25
ii. 25–30
iii. 31–35
iv. 36–40
v. 41–45
vi. Over 45
61. Where were you born? If Other please specify your country (list derived from most common countries of PhD students in Australia).

i. Australia
ii. England
iii. New Zealand
iv. India
v. Italy
vi. Vietnam
vii. Philippines
viii. China
ix. Nepal
x. Malaysia
xi. Brazil
xii. Other (please specify)
62. Is English your first language?

i. Yes
ii. No
63. It is recognised that there are some difficulties for ECRs in working in a research environment in STEMM disciplines. Why do you choose to stay in academia?

### Further Comments

64. Is there anything you would like to add which has not been covered in this survey?

If you wish to, please provide detail about the nature and duration of bullying, and its impact on you, and/or describe any instance of research misconduct which has impacted you here

Open text response (altered question

### Parallel Survey

Opening Comment

This parallel survey invites you to leave contact details if you are interested in taking part in follow up research or receiving results.

### Questions about further contact (parallel survey)

1. Would you like to receive a copy of the final study report? *If so, please leave your email address at Q3*
2. Would you be willing to be contacted by our team for any follow-up research in the future?
3. Please provide your email address below
4. After analysing the data from the survey, we may be conducting interviews to further explore the topics relevant to early-career scientists. Interviews will be conducted in person or via Zoom and will take about one hour. Would you like to be considered for such an interview? Choices: Yes, no, maybe
5. What is your gender? (information required for planning interviews) Choices: Male non-binary female prefer not to say
6. What is the number of years since completion of your highest degree? Choices: 0-4, 5-10, 11-15, other
7. Which of the following best describes your current position within the STEMM research community? By research community, we are referring to all those who conduct or support research.

i. I am a student –[Terminate these]
ii. I am employed / contracted / freelance
iii. I am taking a career-break / on leave (e.g. parental)
iv. I am looking for work / unemployed
v. I am retired
vi. I used to be part of the research community, but no longer am
vii. I have <u>never</u> been part of the research community-[Terminate these]
viii. Other, please specify

#### End Comment

Thank you for taking part in this survey. We may be in touch with you for a follow up interview if we find it necessary and if you have provided your details.

We will keep your details for follow up research, if you have agreed that we may do so.

We will send you research results at the end of the project, if you have asked to receive them.

